# Reversible assembly and disassembly of V-ATPase during the lysosome regeneration cycle

**DOI:** 10.1101/2023.12.22.573005

**Authors:** Ioana Sava, Luther J. Davis, Sally R. Gray, Nicholas A. Bright, J. Paul Luzio

## Abstract

Regulation of the luminal pH of late endocytic compartments in continuously fed mammalian cells is poorly understood. Using normal rat kidney fibroblasts, we investigated the reversible assembly/disassembly of the proton pumping V-ATPase when endolysosomes are formed by kissing and fusion of late endosomes with lysosomes and during the subsequent reformation of lysosomes. We took advantage of previous work showing that sucrosomes formed by the uptake of sucrose are swollen endolysosomes from which lysosomes are reformed after uptake of invertase. Using confocal microscopy and subcellular fractionation of NRK cells stably expressing fluorescently tagged proteins, we found net recruitment of the V1 subcomplex during sucrosome formation and loss during lysosome reformation, with a similar time course to RAB7a loss. Addition of invertase did not alter mTORC1 signalling, suggesting that the regulation of reversible V-ATPase assembly/disassembly in continuously fed cells differs from that in cells subject to amino acid depletion/re-feeding. Using live cell microscopy, we demonstrated recruitment of a fluorescently tagged V1 subunit during endolysosome formation and a dynamic equilibrium and rapid exchange between the cytosolic and membrane bound pools of this subunit. We conclude that reversible V-ATPase assembly/disassembly plays a key role in regulating endolysosomal/lysosomal pH in continuously fed cells.

**Significance statement:**

- In continuously fed cells there is net recruitment of the V1 subcomplex of the proton pumping V-ATPase to endolysosomes as they are formed by kissing and fusion of late endosomes with lysosomes, reducing the luminal pH to promote the activity of lysosomal hydrolases.
- During lysosome reformation, alterations in mTORC1 signalling are not required for the net disassembly of the V-ATPase subcomplex, which occurs with a similar time course to loss of RAB7a.
- Alteration of the dynamic equilibrium and rapid exchange between the cytosolic and endolysosome-bound pools of the V1 subcomplex likely underlies the mechanism of V-ATPase assembly/disassembly.

## Introduction

In mammalian cells, the delivery of endocytosed cargo to lysosomal acid hydrolases is achieved by kissing and fusion events between late endosomes (also known as MVBs, multi-vesicular bodies) and lysosomes (Bright *et al*., 2005; Huotari and Helenius, 2011; Bright *et al*., 2016). These events result in the formation of endolysosomes, which are hybrid organelles with characteristics of both late endosomes and lysosomes and from which re-usable, terminal storage lysosomes are regenerated. This lysosome regeneration cycle (Fig 1A) is analogous to that occurring in the autophagic (i.e. macroautophagic) pathway when lysosomes fuse with autophagosomes to form autolysosomes from which lysosomes are reformed and in the phagocytic pathway in which lysosomes are reformed from phagolysosomes (Yu *et al*., 2010; Bento *et al*., 2016; Yang and Wang, 2021). In all three cases, lysosome reformation occurs after tubulation of the hybrid organelle, followed by scission and maturation events, which are best understood mechanistically in the case of autophagic lysosome reformation (ALR)(Yu *et al*., 2010; Li *et al*., 2016; Chen and Yu, 2017; Yang and Wang, 2021). Formation of endolysosomes creates an acidic compartment, which acts as the principal site of lysosomal acid hydrolase activity within the cell and we have previously shown using live cell microscopy that acid hydrolase activity increases after content mixing between the kissing or fusing organelles, consistent with an increase in acidity. After endocytic lysosome reformation (ELR), the resulting lysosomes are not acid hydrolase active and have a more neutral pH. Variation in luminal pH of lysosomes/endolysosomes at different stages of the regeneration cycle is compatible with older reports that lysosomes within an individual cell exhibit a wide range of pH(Yamashiro and Maxfield, 1987; Butor *et al*., 1995) and with more recent data showing that less acidic lysosomes are preferentially distributed closer to the cell periphery in HeLa cells(Johnson *et al*., 2016), although this latter finding has been challenged(Ponsford *et al*., 2021). The subcellular localization of lysosomes is determined by the balance between the small GTPases RAB7 and ARL8 which interact with kinesin and dynein microtubule motors via different effectors(Jordens *et al*., 2001; Rosa-Ferreira and Munro, 2011; Johnson *et al*., 2016; Jongsma *et al*., 2020), as well as an ER-located ubiquitin ligase system that contributes to their immobilisation in the perinuclear region(Jongsma *et al*., 2016). It has been shown that driving lysosomes to a peripheral location in HeLa cells, by experimentally manipulating the concentration and/or activity of ARL8b, RAB7a effectors and dynein, also results in luminal alkalinization(Johnson *et al*., 2016).

**Figure 1.**
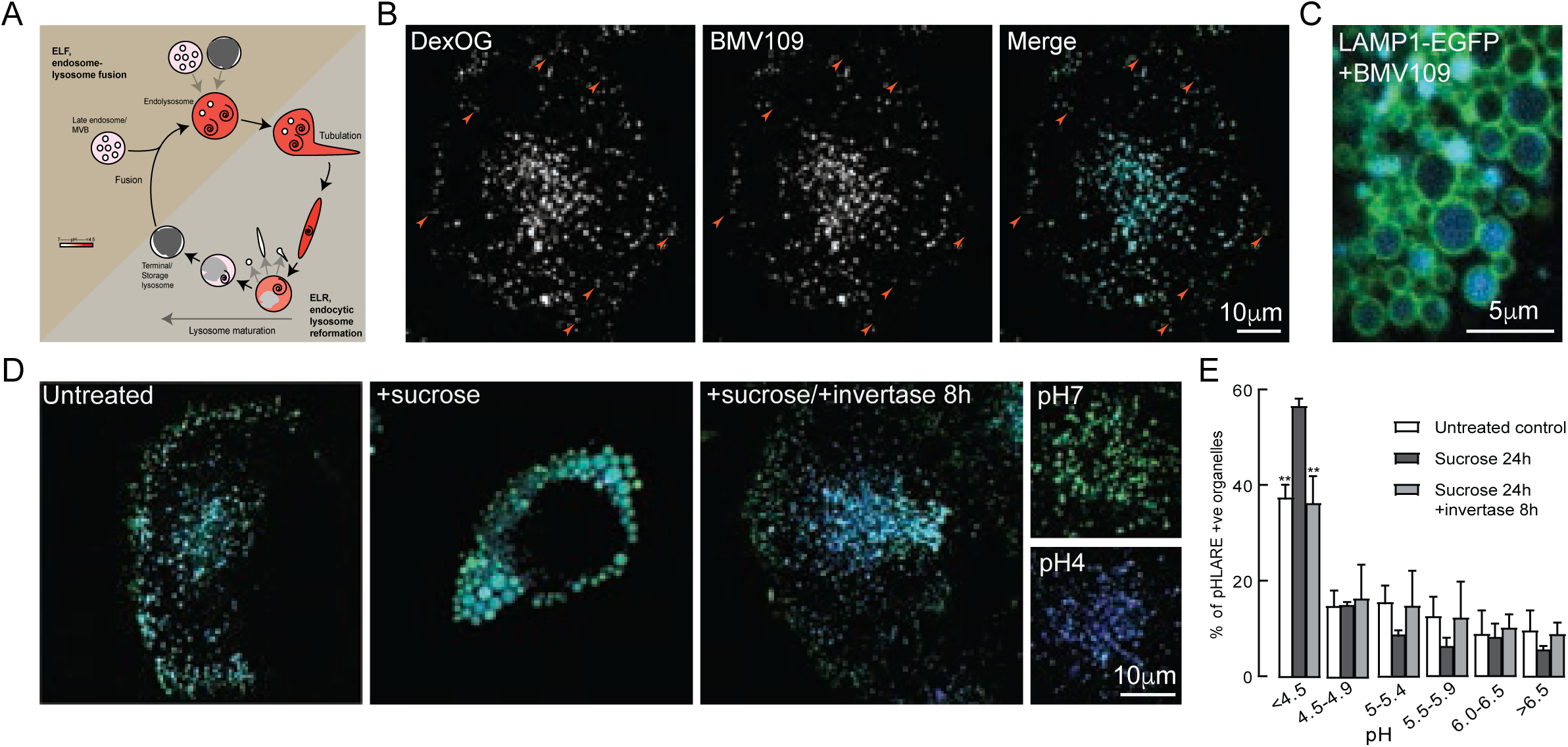
Cathepsin activity and pH of lysosomal organelles in NRK cells. **(A)** Schematic showing the lysosome regeneration cycle, redrawn from (Bright et al., 2016). **(B)** Confocal fluorescence microscopy images of a single representative NRK cell in which terminal endocytic compartments were loaded with DexOG for 4h followed by 20h chase in Dex-OG-free medium and then incubated with BMV109 for 30min. Arrowheads show examples of Dextran-Oregon Green (Dex-OG)-positive/BMV109-negative late endocytic organelles. Colocalization correlation coefficients shown in FigS1A. **(C)** Representative confocal fluorescence microscopy image of part of a live NRK cell stably expressing LAMP1-EGFP and incubated with sucrose for 24h and BMV109 for 30min to show BMV109 labelling of the lumen of sucrosomes. **(D)** Ratiometric imaging of NRK cells loaded (4h pulse/20h chase) with DexOG as the acid-sensitive fluorochrome (green) and Dextran Alexa 647 as an acid insensitive dye (blue), otherwise untreated, incubated with 30mM sucrose for 24h or incubated with sucrose followed by 8h incubation with invertase. Images of parts of cells clamped at pH7 or pH 4 using identical imaging parameters are also shown. **(E)** Distribution of pH in pHLARE-positive organelles in NRK cells stably expressing pHLARE and otherwise untreated, incubated with 30mM sucrose for 24h or incubated with sucrose followed by 8h incubation with 0.5mg/mL invertase. Mean ± SEM of 3 experiments in each of which the pH of individual organelles in 10 cells (single confocal section of each cell) was measured by comparison with a standard curve (Fig S1F) derived from pH clamped cells. **, p < 0.01for difference in % organelles with pH< 4.5 to those in sucrose-treated cells.

The pH of individual endolysosomes and lysosomes is determined by the density and activity of the proton-pumping V-ATPase, counterion transport and passive proton leakage, likely via the TMEM175 channel(Steinberg *et al*., 2010; Mindell, 2012; Ishida *et al*., 2013; Johnson *et al*., 2016; Hu *et al*., 2022; Maxson *et al*., 2022; Zhang *et al*., 2023). An active V-ATPase is essential for the low pH observed in many of these organelles and inhibition with the macrolide antibiotic bafilomycin A1 raises the mean pH of the organelles from 5.3 to 6.5(Rennick *et al*., 2022). Nevertheless, physiological regulation of the lysosomal V-ATPase in mammalian cells remains poorly understood. A mechanism of reversible assembly/disassembly of the V-ATPase V1 and Vo sub-complexes in response to different nutritional conditions has been described in yeast(Parra and Kane, 1998), and increased assembly suggested as the basis of increasing acidification along the endocytic pathway in mammalian cells(Lafourcade *et al*., 2008) as well as the increased acidification of lysosomes during dendritic cell maturation (Trombetta *et al*., 2003). Reversible V-ATPase assembly/disassembly also occurs in mammalian cells in response to amino acid starvation/re-feeding, where it is dependent on mTORC1 activity, such that when mTORC1 is active V1-ATPase subunits reside in the cytosol (Stransky and Forgac, 2015; Ratto *et al*., 2022). It is notable that mTOR activity has also been shown to be necessary for ALR occurring in response to prolonged starvation or exogenous hydrogen peroxide(Yu *et al*., 2010; Zhang *et al*., 2016).

In our present study we have explored the regulation of the V-ATPase during both endosome-lysosome fusion (ELF) and ELR in continuously fed cells by taking advantage of previous work in which we showed that the swollen sucrosomes formed after 24h fluid-phase uptake of sucrose are swollen endolysosomes which have recruited all of the terminal storage lysosomes(Bright *et al*., 1997; Bright *et al*., 2016). Subsequent fluid phase uptake of invertase initiates ELR, approximately synchronously, from all the sucrosomes. We have investigated whether V-ATPase assembly increases during ELF and if disassembly occurs during ELR. We have also examined whether disassembly is linked to the loss of RAB7 and whether changes in mTORC1 activity are required for alterations in the V-ATPase during ELR in continuously fed cells.

## Results

Throughout our experiments, we have used NRK (normal rat kidney) fibroblasts, in which sucrosomes may be generated and lysosomes reformed, without activation of autophagy (Bright *et al*., 2016). For many experiments we have used NRK cells transduced with retroviral vectors containing weak promoters to achieve stable expression of tagged proteins at mild levels of over-expression (Supplementary Table S1).

### Sucrosomes are swollen endolysosomes containing active cathepsins in acidic lumens and from which less acidic lysosomes can be reformed

To study the localization of active cathepsins we used the cell-permeable pan-cathepsin activity-based probe BMV109, which is fluorescent after it covalently binds to cysteine cathepsins(Withana *et al*., 2016; Repnik *et al*., 2017), as well as membrane-permeable Magic Red cathepsin substrates, which liberate fluorescent cresyl violet after hydrolysis(Creasy *et al*., 2007; Bright *et al*., 2016). Using BMV109 avoided the problem of liberated cresyl violet potentially migrating to an acidic compartment other than its site of generation (Ostrowski *et al*., 2016) and confirmed that in untreated NRK cells there is a population of late endocytic compartments observed by fluorescent dextran uptake that is cathepsin inactive (Fig 1B, S1A), consistent with this population being reusable terminal lysosomes (Bright *et al*., 2016). When NRK cells stably expressing LAMP1-EGFP were incubated with sucrose to form sucrosomes and treated with BMV109, the sucrosome lumens became fluorescent confirming that they contained active acid hydrolases (Fig 1C, S1B), as previously shown with Magic Red cathepsin substrates and by acid phosphatase staining in EM (Bright 2016). Because the sucrosomes are swollen organelles it was possible, using confocal microscopy, to differentiate the luminal BMV109 fluorescence from the LAMP1-EGFP fluorescence on the limiting membrane, which was not achievable for the much smaller endolysosomes and lysosomes in control cells. Thus, we could monitor formation of sucrosomes and their collapse after invertase uptake (Fig 1C and S1B) by measuring the colocalization (Pearson’s correlation coefficient) of BMV109 with LAMP1-EGFP fluorescence (Fig S1C). After invertase uptake, the Pearson’s coefficient returned to that in untreated cells by 2h, reflecting shrinkage of the sucrosomes rather than their complete loss, which was previously observed by EM in NRK cells (Fig S1D) (Bright *et al*., 1997), to be complete by ∼4 h (Fig S1D) (Bright *et al*., 1997).

To visualise the variation in pH of individual lysosomes, endolysosomes and sucrosomes, we used dual fluorochrome imaging after endocytic uptake of pH-sensitive dextran Oregon green (DexOG) together with pH-insensitive dextran Alexa (DexA) 647. In control, untreated NRK cells, less acidic organelles (green in Fig 1D) were enriched at the cell periphery and more acidic organelles (cyan, turquoise and blue) enriched in the juxtanuclear region, as had previously been observed in HeLa cells (Johnson *et al*., 2016). When sucrosomes were formed, they were different shades of cyan and turquoise (Fig1D), consistent with them being cathepsin active(Bright *et al*., 2016)(FigS1A ie BMV pictures). After uptake of invertase for 8h, the peripheral, neutral lysosomes (green) had reappeared (Fig1D). Using cells in which the genetically encoded pH probe pHLARE(Webb *et al*., 2021) was stably expressed, a shift to a higher proportion of organelles with pH <4.5 was observed after formation of sucrosomes, which was reversed after invertase uptake (Fig 1E). The lysosomal/endosomal compartment in control, untreated NRK cells expressing the probe had a mean pH of 5.18±0.04, in cells incubated with sucrose for 24h, 4.60±0.16 and in cells after subsequent 8h incubation with invertase, 5.11±0.12 (Fig S1E).

In an initial experiment to test whether assembly/disassembly of V-ATPase may be responsible for the variation in pH observed, we generated a mixed cell population stably expressing EGFP-tagged V1G1, a ubiquitous isoform of the V1ATPase subunit G which is a peripheral stalk protein in the V-ATPase proton pump (Table S1). A punctate distribution of V1G1-EGFP was observed as well as a cytosolic pool and the puncta colocalized well with Magic Red staining of cathepsin B activity (Fig S1G). In addition, there was partial overlap of V1G1-EGFP-positive puncta with antibody staining of the endogenous Vo subunit Voa3 (TCIRG1), an isoform concentrated in late endosomes and lysosomes (Toyomura *et al*., 2003)(Fig S1H). It was noticeable that in the cell periphery there were many Voa3-positive puncta that were V1G1-EGFP-negative (Fig S1H). Whilst this was consistent with V-ATPase disassembly being associated with an enrichment of alkalinized lysosomes towards the cell periphery, the variation in expression of V1G1-EGFP in different cells of the mixed population guided us towards studying the dynamics of V-ATPase subunits in clonal cell lines (Table S1).

### V1 and Vo subunits of V-ATPase enter reformation tubules as ELR takes place from sucrosomes after addition of invertase

In clonal cell lines stably expressing either V1G1-EGFP or Voa3-EGFP and incubated with sucrose, clusters of sucrosomes were observed in which the EGFP-tagged proteins were localised to the rims of the organelles, the lumens of which were filled by cresyl violet when incubated with Magic Red cathepsin substrates (Fig 2A, B). Following uptake of invertase, reformation tubules were formed as seen previously (Bright *et al*., 2016)and these were labelled with both the EGFP-tagged V-ATPase subunits when observed by live cell imaging in a confocal microscope (Fig 2C, D and Video 1)). Confirmation that V1G1-EGFP was entering the tubules was obtained by correlative light and electron microscopy (CLEM) using thick sections from unroofed cells, which allowed more efficient immunolabelling than conventional immuno-EM on frozen thin sections (Fig 2E and S2A,B). Thus, disassembly of the V-ATPase does not occur as a reformation tubule is generated but could take place after scission as the reforming lysosome matures.

**Figure 2.**
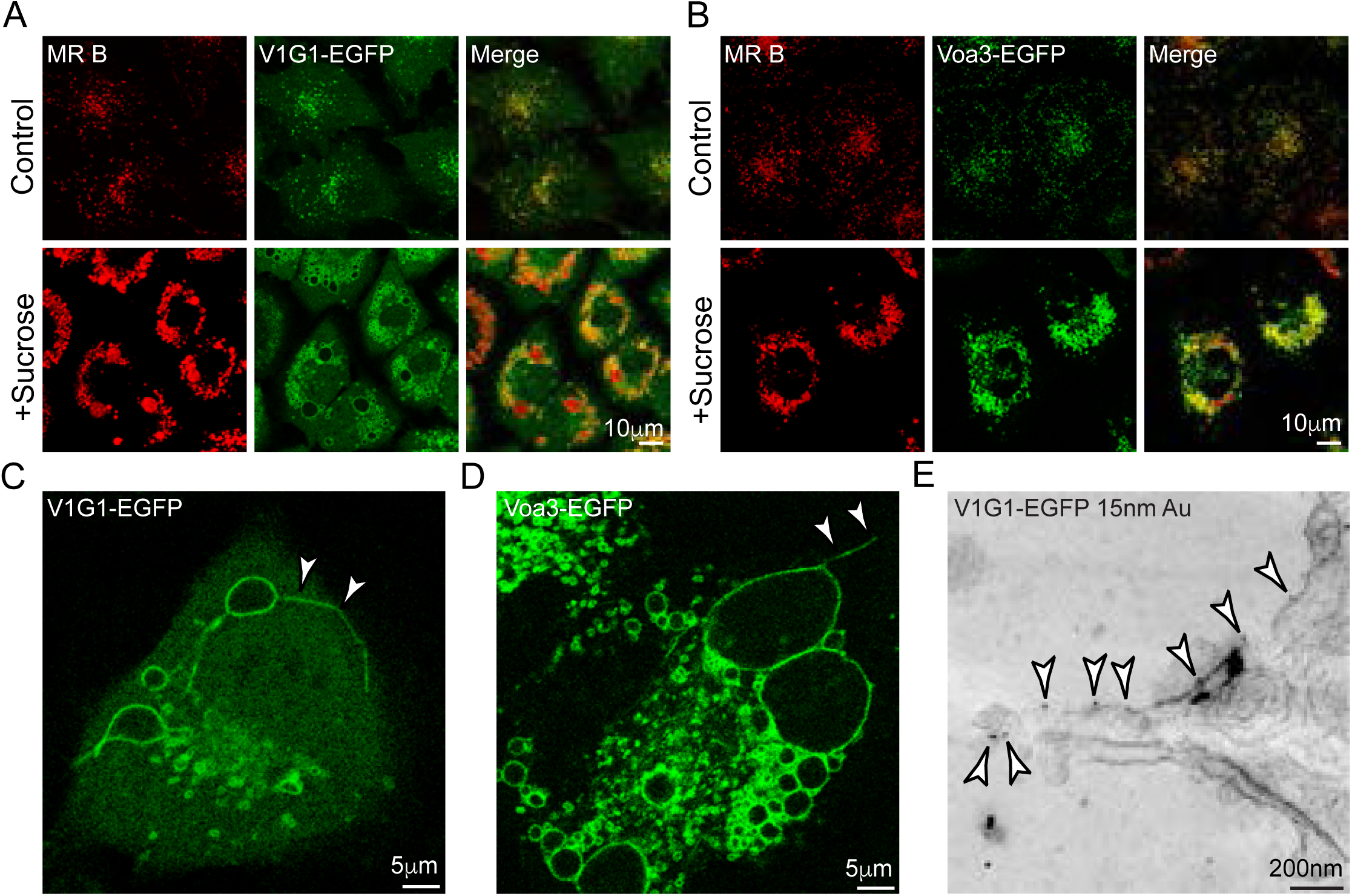
Localization of EGFP-tagged V-ATPase proteins in stably expressing NRK cells. **(A, B)** Confocal fluorescence images of untreated (control) clonal cells expressing V1G1-EGFP (A) or Voa3-EGFP (B) or cells incubated with 30mM sucrose for 24h were incubated with Magic Red cathepsin B substrate (MR B) for 2min prior to imaging. **(C)** Still from Video1 showing V1G1-EGFP entering lysosome reformation tubule (arrowheads) after incubating V1G1-EGFP cells containing sucrosomes with 0.5 mg/mL invertase for 1h. **(D)** Confocal image showing Voa3-EGFP entering lysosome reformation tubule (arrowheads) after incubating Voa3-EGFP cells containing sucrosomes with invertase as in (C). **(E)** Section of immunoelectron micrograph of unroofed V1G1-EGFP cell shown in Suppl Fig S2B following unroofing and CLEM. Arrowheads indicate gold particles identifying V1G1-EGFP in lysosome reformation tubule forming from the edge of a sucrosome.

EM of unroofed cells also revealed that the clustered appearance of sucrosomes seen by fluorescence microscopy was due to filamentous tethers between the organelles (Fig S2C). Immunolocalization of the HOPS protein VPS18 by immunofluorescence microscopy showed its presence on sucrosomes and immuno-EM of unroofed cells showed sparse labelling close to/on the tethers suggesting that the tethering is mediated by the HOPS complex (FigS2D). The sparse labelling may be due to poor antibody access to epitopes in tightly packed tether arrays as previously discussed(Davis *et al*., 2021).

### Confocal microscopy demonstrated increased colocalization of RAB7a, RILP and the V1G1 subunit of V-ATPase during sucrosome formation and a decrease during lysosome reformation

To test if there was increased recruitment of the V-ATPase V1 subcomplex during sucrosome formation and loss during lysosome reformation and their relationship to the presence of RAB7a and RILP, we studied the colocalization of tagged proteins using confocal microscopy. Stably expressing mixed cell populations were generated by separately transducing a clonal cell line expressing LAMP1-mCherry with retroviral vectors encoding each of EGFP-tagged Voa3, V1G1, RAB7a, and ARL8b (Table S1, FigS2E). To avoid excessive perinuclear aggregation of endolysosomal organelles caused by overexpression of RILP ((Cantalupo *et al*., 2001; Jordens *et al*., 2001), a clonal line of cells expressing LAMP1-mCherry and EGFP-RILP was generated and experiments were conducted after ∼85% depletion of endogenous RILP using siRNA (Fig S2E, F). Colocalization of each EGFP-tagged construct with LAMP-mCherry was assessed by calculating Manders’ and Pearson’s coefficients in control, untreated NRK cells, cells after formation of sucrosomes and cells at different time points after incubation with invertase to initiate reformation of lysosomes (Fig 3A-F and S3A-E). Colocalization of the EGFP-tagged integral membrane protein Voa3 and of ARL8b-EGFP with LAMP1-mCherry varied little throughout the experiment (Fig. 3F). However, colocalization of the EGFP-tagged V1G1, RAB7a and RILP with LAMP1-mCherry increased when sucrosomes were formed and decreased after addition of invertase, with statistically significant decreases being observed at 4-8 hours (Fig 3B-D,F and S3B-D). These data are consistent with reversible assembly/disassembly of the V-ATPase being associated respectively with gain/loss of RAB7a and RILP.

**Figure 3.**
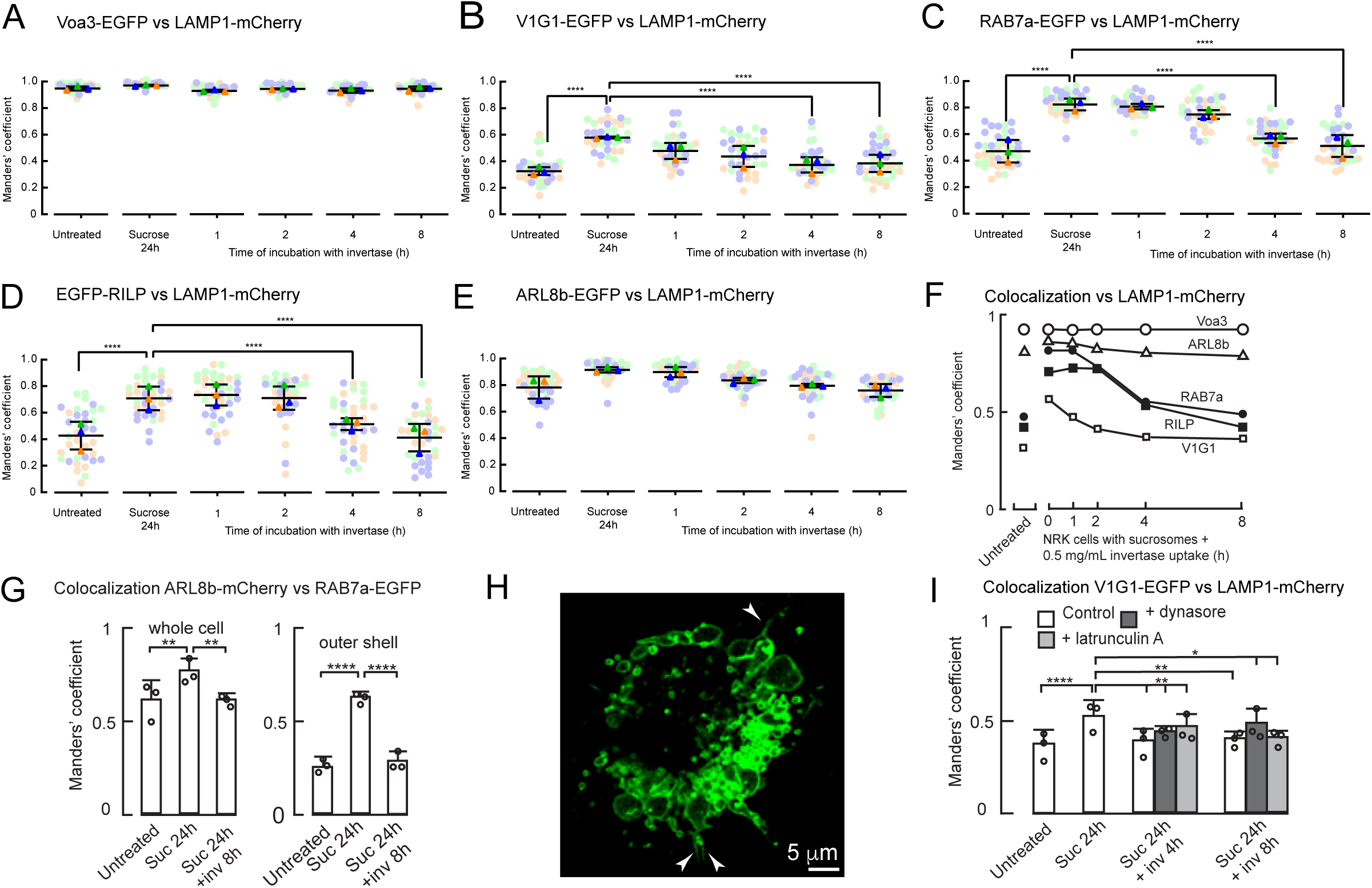
Colocalization of EGFP-tagged proteins with LAMP1-mCherry or ARL8b-mCherry in stably expressing NRK cells. **(A-E)** Manders’ colocalization coefficients for EGFP-tagged proteins vs LAMP1-mCherry in untreated cells, cells incubated with 30mM sucrose for 24h or incubated with sucrose followed by 1-8h incubation with 0.5mg/mL invertase. Mixed stable cells used except in (D) in which a clonal LAMP1-mCherry/ RILP-EGFP cell line, 86.7±1.9% (3) depleted of endogenous RILP by siRNA-mediated knockdown, was used. **(F)** Summary of changes in mean Manders’ colocalization coefficients shown in panels A-E. **(G)** Manders’ colocalization coefficients (whole cell or outer shell) for ARL8b-mCherry vs RAB7a-EGFP in untreated mixed stable ARL8b-mCherry/ RAB7a-EGFP cells, cells incubated with 30mM sucrose for 24h or incubated with sucrose followed by 8h incubation with 0.5mg/mL invertase. **(H)** Confocal image showing RAB7a-EGFP entering lysosome reformation tubules (arrowheads) after incubating RAB7a-EGFP cells containing sucrosomes with 0.5 mg/mL invertase for 1h **(I)** Manders’ colocalization coefficients for V1G1-EGFP vs LAMP1-mCherry in the presence or absence of 80μM dynasore or 1μM latrunculin A in otherwise untreated LAMP1-mCherry/ V1G1-EGFP mixed stable cells, cells incubated with 30mM sucrose for 24h or incubated with sucrose followed by 8h incubation with 0.5mg/mL invertase. In panels A-E, G and I, mean ± SEM of 3 experiments (minimum 10 cells per experiment) shown. *, p<0.05; **, p<0.01, ***, p<0.005; ****, p<0.001.

To study the colocalization of RAB7a with ARL8b, we generated a mixed cell population stably expressing EGFP tagged RAB7a and mCherry-tagged ARL8b (Fig S2E). We observed statistically significant changes in the colocalization coefficients for these two small GTPases during sucrosome formation and after 8h of lysosome reformation in these cells, which were greater for organelles in the outer ‘shell’ of the cytoplasm (Fig 3G and S3F,G). The data were consistent with a reduced RAB7a density and an enrichment of ARL8b relative to RAB7a on peripheral lysosomes as observed previously in HeLa cells(Johnson *et al*., 2016), as well as the recruitment of these peripheral lysosomes into the sucrosomes and their reformation from the sucrosomes, although we did not pursue this further.

As was the case with V1G1-EGFP, RAB7a-EGFP also entered reformation tubules after invertase addition to cells containing sucrosomes (Fig 3H), suggesting that its loss also occurs after scission as the reforming lysosome matures. Interestingly, the addition of the dynamin inhibitor dynasore (Macia *et al*., 2006), which has been shown to block dynamin-mediated cleavage of reformation tubules in ALR (Schulze *et al*., 2013), did not prevent the decrease in colocalization of V1G1-EGFP with LAMP1-mCherry after delivery of invertase to sucrosomes (Fig 3I and S3H,I), although transmission EM showed the appearance of ‘daisy chains’ of swollen organelles with interconnected lumens (Fig S3J), consistent with the inhibition of fission of the reformation tubules, which we still observed forming and retracting in live cell microscopy (Video 2). These colocalization date suggest that cleavage of reformation tubules is not necessary for the disassembly of the V-ATPase, even though disassembly most likely occurs post-cleavage in untreated cells. We also saw no effect of latrunculin A, which prevents actin polymerisation, on the decrease of the colocalization of V1G1-EGFP with LAMP1-mCherry after delivery of invertase to sucrosomes (Fig 3I and S3H,I), despite actin polymerisation being implicated in V-ATPase sorting in *Dictyostelium* (Carnell *et al*., 2011).

### Biochemical evidence that the V1 subcomplex of V-ATPase is recruited during sucrosome formation and lost during lysosome reformation

The data from the microscopy colocalization experiments were consistent with reversible assembly and disassembly of V-ATPase during sucrosome formation and lysosome reformation, but to measure this directly we carried out subcellular fractionation and immunoblotting of the LAMP1-mCherry/V1G1-EGFP cells. Immunoblotting of membrane fractions has previously been used to show the role of amino acid availability in modulating V-ATPase assembly (Stransky and Forgac, 2015), so we prepared 100,000g membrane pellets from control, untreated cells, cells with sucrosomes and cells in which lysosome reformation was occurring after invertase uptake. SDS-polyacrylamide gel electrophoresis (SDS-PAGE) and immunoblotting under conditions in which the loading of LAMP1-mCherry was equal for each membrane pellet, showed that the ratio of the Vo subunit Vod1 to LAMP1-mCherry was constant in each membrane pellet, but the ratio of V1G1-EGFP to Vod1 increased in membrane pellets from cells containing sucrosomes and decreased back to untreated cell levels after 8h incubation with invertase (Fig 4A,B). These data strongly supported the hypothesis that there was reversible assembly and disassembly of V-ATPase during sucrosome formation and lysosome reformation. However, there was considerable variability between experiments and the 100,000g membrane pellets contain membranes from many organelles other than endolysosomes/sucrosomes and lysosomes.

**Figure 4.**
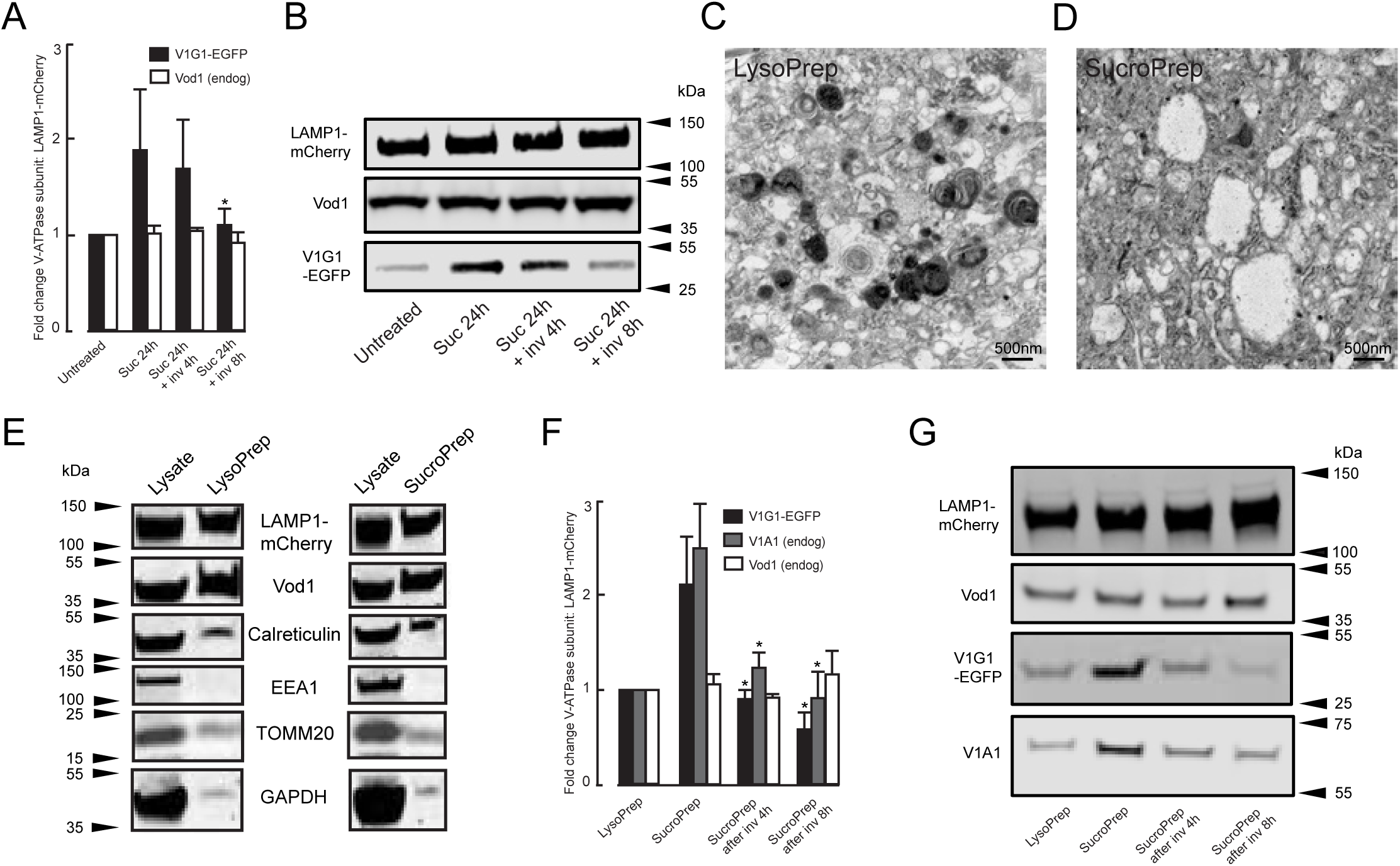
V-ATPase assembly/disassembly during sucrosome formation and lysosome reformation. **(A)** Change in V-ATPase subunit concentration relative to LAMP1-mCherry in 100,000g membrane pellets from untreated LAMP1-mCherry/ V1G1-EGFP mixed stable cells, cells incubated with 30mM sucrose for 24h or incubated with sucrose followed by 4 or 8h incubation with 0.5mg/mL invertase. Mean±SEM of 6 experiments. *, p<0.05 relative to membrane pellet from cells incubated with sucrose alone. **(B)** Immunoblots from a single representative experiment of those summarised in (A). **(C, D)** Transmission electron micrographs of magnetically isolated lysosomes (LysoPrep, C) and sucrosomes (SucroPrep, D). **(E)** Representative immunoblots of selected markers showing enrichment of LAMP1-mCherry and Vod1 relative to other subcellular markers in a LysoPrep and a SucroPrep from LAMP1-mCherry/V1G1-EGFP cells compared to cell lysates. Gel track loading: 15 L/1.5mL total lysate; LysoPrep and SucroPrep 15 L/0.3mL total preparation. **(F)** Change in V-ATPase subunit concentration relative to LAMP1-mCherry in Lysopreps from untreated LAMP1-mCherry/V1G1-EGFP cells and Sucropreps from cells incubated with 30mM sucrose for 24h or incubated with sucrose followed by 4 or 8h incubation with 0.5mg/mL invertase. Mean±SEM of 3 experiments. *, p<0.05 relative to SucroPrep from cells incubated with sucrose alone. **(G)** Immunoblots from a single representative experiment of those summarised in (F).

To quantify the assembly/disassembly of V-ATPase on endolysosomes/sucrosomes and lysosomes more directly we modified previously published rapid magnetic isolation procedures (Rofe and Pryor, 2016; Tharkeshwar *et al*., 2020; Prentzell *et al*., 2021), following uptake of magnetic dextran (DexoMAG, Mr 10,000) into our NRK cells. Our well characterised protocol for accumulating fluorescent dextran (Mr 10,000) and gold labelled bovine serum albumin in NRK lysosomes and endolysosomes after fluid phase endocytosis for 4h followed by a 20 h chase (Bright *et al*., 1997; Bright *et al*., 2005; Bright *et al*., 2016) had to be modified because significant numbers of cells detached from the substrate after uptake of DexoMAG with this protocol. A shorter loading 1h uptake/2h chase protocol was therefore used after we confirmed, using fluorescent dextrans, that this resulted in the same intracellular colocalization with preloaded dextran (4h uptake/20h chase) as a 4h uptake/24h chase protocol (Fig S4A). Using rapid magnetic isolation, we were able to generate enriched fractions of endolysosomes/lysosomes (LysoPrep) from the untreated LAMP1-mCherry/V1G1-EGFP cells and also sucrosomes (SucroPrep) following uptake of sucrose. Transmission EM showed that the LysoPreps were enriched with electron dense lysosomes (Fig 4C), which were labelled with anti-LAMP1 in immuno-EM (Fig S4B, whereas the SucroPreps had few, if any, electron dense lysosomes and were characterised by the presence of swollen electron lucent organelles (Fig 4D), which were labelled on their limiting membrane with anti-LAMP1 in immuno-EM (Fig S4C). SDS-PAGE and immunoblotting showed that the LysoPrep and SucroPrep fractions showed a high yield of LAMP1-mCherry-positive (>15%) and Vod1-positive membranes and were enriched relative to a range of markers for non-lysosomal membranes, other organelles and cytosol (Fig 4E, S4D,C). Further SDS-PAGE and immunoblotting of LysoPreps and SucroPreps from untreated cells, cells with sucrosomes and sucrososome-containing cells incubated with invertase for 4h, under conditions in which the loading of LAMP1-mCherry was equal for each, showed that whereas the ratio of the endogenous Vo subunit Vod1 to LAMP1-mcherry was constant in each preparation, the ratio of V1G1-EGFP and endogenous V1A1 to LAMP1-mCherry increased in the SucroPreps and decreased back towards control cell LysoPrep levels after incubation with invertase for 4 or 8h (Fig 4F,G). The data were less variable between experiments than the data from the 100,00g pellets and are consistent with reversible assembly and disassembly of V-ATPase during sucrosome formation and lysosome reformation.

### VIG1-EGFP is recruited to the limiting membrane of endolysosomes during endosome-lysosome kissing and fusion

The data from the colocalization microscopy and immunoblotting experiments described above, on cells incubated with sucrose, showed an increase in association of the V-ATPase V1 subcomplex with LAMP1/Vo subcomplex-containing membranes when sucrosomes were formed. Although there is strong evidence that sucrosomes are swollen endolysosomes, we decided to use live cell fluorescence confocal microscopy to see whether V1G1-EGFP is also recruited to newly formed endolysosomes in NRK cells that were not incubated with sucrose. Terminal lysosomes in NRK cells stably expressing V1G1-EGFP, were loaded with DexA647 and late endosomes with DexA546 as previously described(Bright *et al*., 2005). We searched for kissing and fusion events between these organelles, observed by content mixing. Following the commencement of content mixing, we observed the gradual recruitment of V1G1-EGFP to the peripheral membrane of the newly formed endolysosome (Video 3, Fig 5A, B, S5A-G). Usually, there was a lag between the commencement of content mixing and the recruitment of V1G1-EGFP (Fig 5A, B, S5H), consistent with the lag previously observed before the appearance of cresyl violet in a newly formed endolysosome, when NRK cells where incubated with cathepsin B MR substrate in the surrounding medium(Bright *et al*., 2016). These data taken together are consistent with assembly of the V-ATPase preceding acidification and activation of cathepsins when endolysosomes are formed.

**Figure 5.**
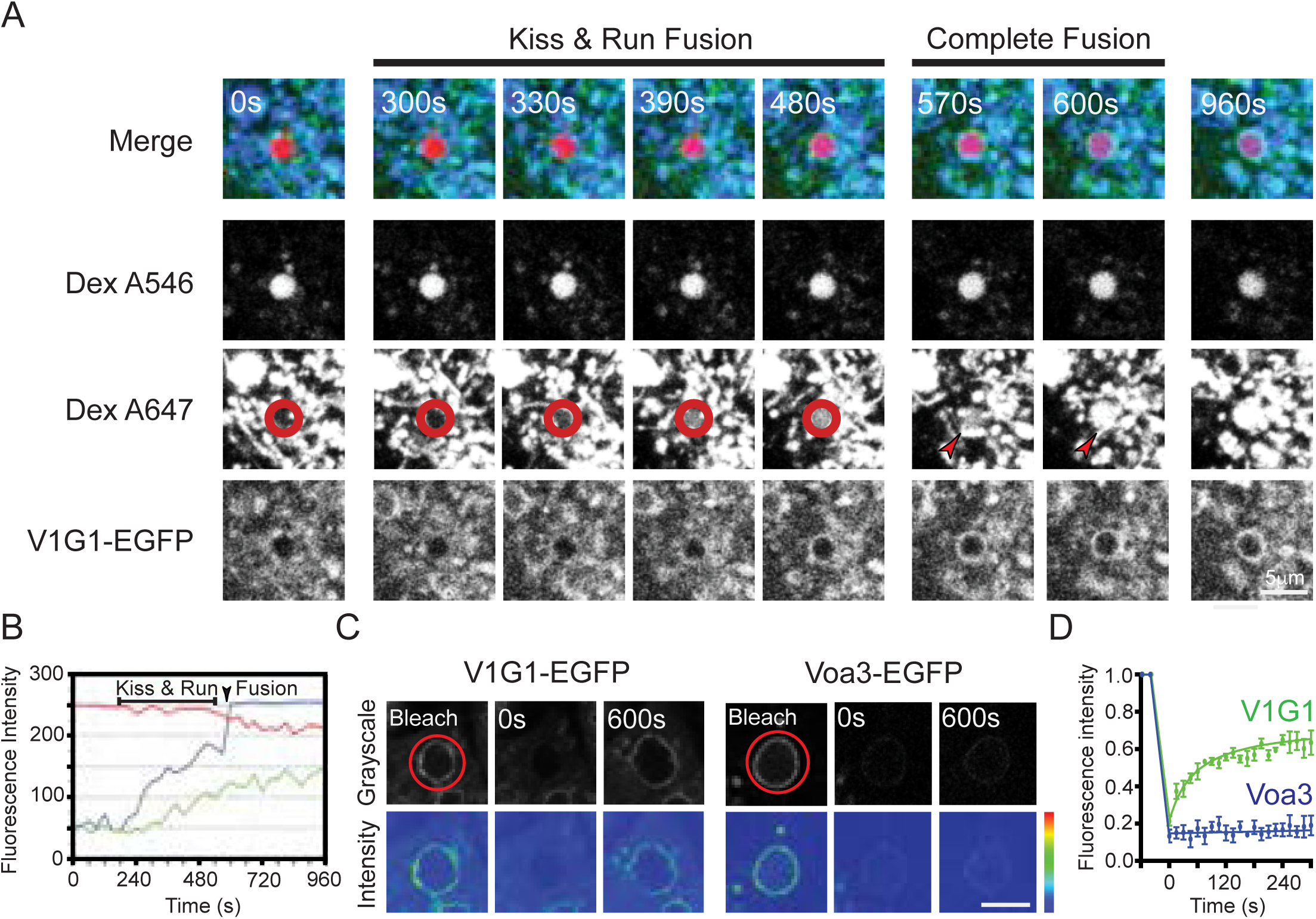
The dynamics of V-ATPase V1 subunit recruitment to endolysosomes in live cells. **(A)** Terminal endocytic compartments of clonal NRK cell line expressing V1G1-EGFP were pre-loaded with Dex A647 for 4h followed by a 20h chase in Dex A-free medium, and then late endosomes were loaded with Dex A546 by uptake for 10min followed by a 5min chase in Dex A546-free medium. Selected images from time-lapse confocal microscopy (video 3) of the living cells for 16min, showing that kissing and fusion of a Dex A546-positive late endosome with Dex A647-positive terminal endocytic compartments (lysosomes) resulted in the gradual acquisition of Dex A647 (blue) in the Dex A546-positive organelle (red and red circle) and a subsequent rise in recruitment of V1G1-EGFP (green) to the limiting membrane of the resulting endolysosome. Arrowheads indicate the organelles that fuse. **(B)** Mean grayscale intensity profile of Dex A546 (red), Dex A647 (blue) and V1G1-EGFP (green) over 16min for the organelle outlined in the red circle in (A) and shown in Video3. **(C)** Representative FRAP images from time-lapse confocal microscopy (Video 4) of living clonal V1G1-EGFP cells and clonal Voa3-EGFP cells showing fluorescence bleaching of a sucrosome from each cell type, followed by fluorescence recovery of V1G1-EGFP but not Voa3-EGFP. Images shown in grayscale or as an intensity map (high intensity, red; mid intensity, green; low intensity, blue). Bar = 5 μm. **(D)** Alteration in fluorescence intensity (grayscale, mean±SEM) in FRAP experiments on 3 sucrosomes in clonal V1G1-EGFP cells and on 3 sucrosomes in clonal Voa3-EGFP cells.

### The dynamic nature of V1G1 association with the sucrosome limiting membrane

The data above demonstrate net assembly of V-ATPases when endolysosomes, including sucrosomes, are formed and disassembly when lysosomes are reformed. However, the time courses of our experiments are much longer than a previous report of the half time of dynamic exchange of V1-ATPase subunits bound to presynaptic vesicles in hippocampal neurons (Bodzeta *et al*., 2017). We therefore examined whether V1G1-EGFP is dynamically associated with the sucrosome membrane, using fluorescence recovery after photobleaching (FRAP) experiments. As was previously observed for GFP-tagged subunits on synaptic vesicles, the fluorescence signal from the V1G1-EGFP labelled sucrosomes rapidly recovered after photobleaching with, in this case, a t1/2 of ∼45s, suggesting that the pool of V1G1-EGFP on the membrane is in dynamic equilibrium with the cytosolic pool and rapidly exchanging with it (Video 4, Fig 5C, D). As predicted, no recovery of fluorescence occurred after photobleaching of Voa3-EGFP on the sucrosome membrane (Video 4, Fig 5C, D), unless there was subsequent fusion with another Voa3-EGFP-positive vesicle or organelle (data not shown). Similar results were obtained after invertase uptake into the sucrosomes, although the transient nature of lysosome reformation tubules made it difficult to assess whether loss to/recruitment of V1G1-EGFP from the cytosolic pool continued to occur on reformation tubules before they detached from the parent sucrosome (data not shown).

### Changes in mTOR signalling are not required for the assembly and disassembly of the V-ATPase during sucrosome formation and lysosome reformation in the NRK cells

Reactivation of mTORC1 signalling has been described as necessary for ALR after prolonged serum starvation of NRK cells(Yu *et al*., 2010), so we investigated whether there was any change in mTORC1 signalling during the initiation of reformation of lysosomes from sucrosomes. mTORC1 signalling was evaluated by SDS-PAGE and immunoblotting of the phosphorylated forms of the eukaryotic initiation factor 4E-binding protein 4E-BP1, the p70 S6 kinase 1 (P70S6K) and its target ribosomal protein S6, a component of the 40S ribosomal subunit(Klionsky *et al*., 2021). Although, sucrose has been reported to activate autophagy via mTOR-dependent and -independent pathways (Chen *et al*., 2016; Khan *et al*., 2017) in other cell types, we observed no significant effect of 30mM sucrose on the amount of these phosphorylated proteins in the NRK cells after 30min or 24h incubation and no effect of invertase when added to the sucrosome-containing cells for 30min or 1h (Fig 6A, B). We did see a variable and non-significant reduction in phosphorylation after 30 min treatment, but did not investigate this further because it was not apparent after 24h, the starting point for our lysosome reformation experiments. As expected, the mTOR inhibitor torin greatly reduced the amount of the three phosphorylated proteins (Fig 6A,B, S6A). Consistent with changes in mTOR signalling not being required for the assembly and disassembly of the V-ATPase during sucrosome formation and lysosome reformation in the NRK cells, the presence of torin had no effect on either the increase in colocalization of V1G1-EGFP with LAMP-mCherry during sucrosome formation or the time course of its decrease after incubating the sucrosome-containing cells with invertase (Fig 6C, S6B). Torin also had no effect on the assembly or disassembly of the V-ATPase, when assessed by immunoblotting of magnetically isolated endolysosomal organelles from cells incubated with sucrose or sucrose followed by invertase (Fig 6D,E). We also observed no effect of torin on the mean pH of the endolysosomal/lysosomal compartments nor any significant shift in the proportion of higher or lower pH in NRK cells stably expressing the pH probe pHLARE (Fig 6F, S6C).

**Figure 6.**
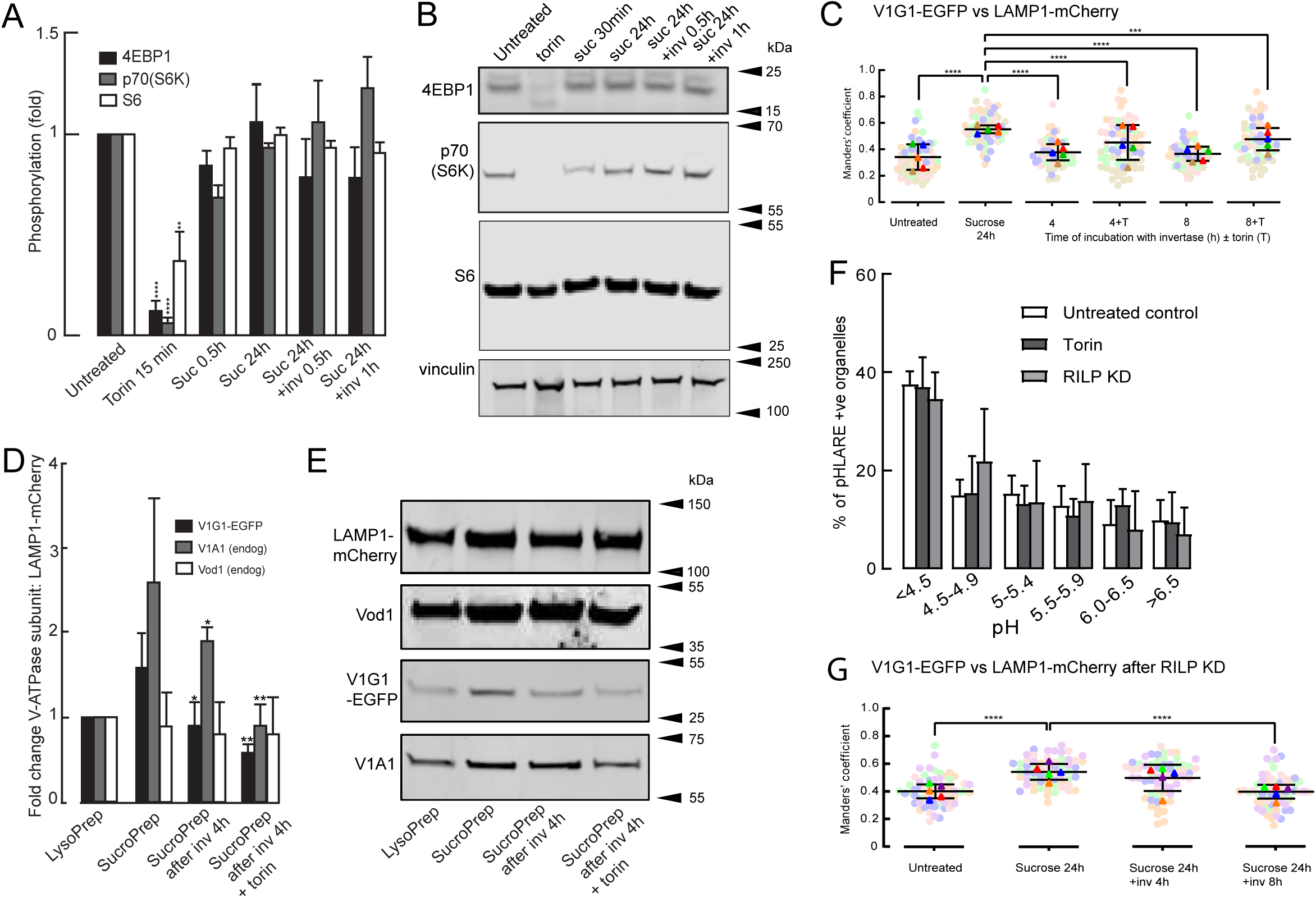
mTORC1 signalling, RILP and V-ATPase assembly/disassembly in continuously fed cells. **(A)** Alteration of phosphorylation of mTORC1 substrates in NRK cells treated with 1mM torin or incubated with 30 mM sucrose for 30 min or 24h ± subsequent incubation with 0.5mg/mL invertase for 30min or 1h. Treatment with 1 μM torin for 15 min used as a control. Data are mean±SEM of 3 experiments. **, p<0.01, ****, p<0.001 relative to untreated cells. **(B)** Immunoblots of phosphorylated mTORC1 substrates from a single representative experiment of those summarised in (A). Vinculin used as loading control. **(C)** Manders’ colocalization coefficients for V1G1-EGFP vs LAMP1-mCherry in otherwise untreated LAMP1-mCherry/ V1G1-EGFP mixed stable cells, cells incubated with 30mM sucrose for 24h or incubated with sucrose followed by 4 or 8h incubation with 0.5mg/mL invertase ± 0.25 μM torin. Mean ± SEM of 3 experiments (minimum 10 cells per experiment) shown. ***, p<0.005; ****, p<0.001. **(D)** Change in V-ATPase subunit concentration relative to LAMP1-mCherry in Lysopreps from untreated LAMP1-mCherry/V1G1-EGFP cells and Sucropreps from cells incubated with 30mM sucrose for 24h or incubated with sucrose followed by 4h incubation with 0.5mg/mL invertase ± 0.25 μM torin. Mean±SEM of 3 experiments. *, p<0.05; **, p<0.01 relative to SucroPrep from cells incubated with sucrose alone. **(E)** Immunoblots from a single representative experiment of those summarised in (D). **(F)** Distribution of pH in pHLARE-positive organelles in NRK cells stably expressing pHLARE and otherwise untreated, incubated with 0.25 μM torin 15min or following siRNA-mediated knockdown of RILP (96.1±1.3% (3) depletion). Mean ± SEM of 3 experiments in each of which the pH of individual organelles in 10 cells (single confocal section of each cell) was measured by comparison with a standard curve (Fig S1F) derived from pH clamped cells. **(G)** Manders’ colocalization coefficients for V1G1-EGFP vs LAMP1-mCherry in LAMP1-mCherry/ V1G1-EGFP mixed stable cells following siRNA-mediated knockdown of RILP (85.1±2% (5) depletion) and otherwise untreated, incubated with 30mM sucrose for 24h or incubated with sucrose followed by 4 or 8h incubation with 0.5mg/mL invertase. Mean ± SEM of 5 experiments (minimum 10 cells per experiment) shown. ****, p<0.001.

### Is RILP binding to V1G1 necessary and sufficient for the assembly and disassembly of the V-ATPase during sucrosome formation and lysosome reformation ?

The rapid, continuous exchange of a V1 subunit between a membrane-bound and cytosolic pool observed in our FRAP experiments described above implies that a mechanism that promotes an increase in V-ATPase assembly would, if switched off or lost, result in net disassembly. Such a mechanism could simply involve the binding of RILP to V1G1 (De Luca *et al*., 2014), which has been suggested to explain the pH difference of juxtanuclear and peripheral lysosomes (Johnson *et al*., 2016). However, when we depleted RILP by >90% in NRK cells stably expressing the pHLARE probe, we saw no change in the mean pH of the endolysosomal/ lysosomal compartment nor any significant shift in the proportion of higher or lower pH organelles (Fig 6F, S6C). Moreover, depletion of RILP by ∼85% in LAMP1-mCherry/V1G1-EGFP cells resulted in no difference in the changes in colocalization of V1G1-EGFP with LAMP1-mCherry during formation of sucrosomes and reformation of lysosomes, after 8h incubation with invertase, compared with undepleted cells (Fig6G, S6D, E). The siRNA-mediated depletion is not a full knockout and in our lysosome reformation experiments we did not observe a statistically significant reduction in colocalization of V1G1-EGFP with LAMP1-mCherry after 4h incubation with invertase (Fig6G, S6D), which always occurred in experiments using cells with a full complement of RILP (Fig3B, S3B). Nevertheless, these data suggest it is most unlikely that the presence of RILP is sufficient to explain fully the assembly/disassembly of the endolysosomal V-ATPase during ELF/ELR in continuously fed cells.

## Discussion

Our study has provided evidence that regulation of the V-ATPase, responsible for the luminal acidification of endolysosomal organelles in continuously fed cells is mediated by the reversible assembly and disassembly of the V1 and Vo sub-complexes and that this is not linked to a change in activity of mTORC1. Our data add to previous evidence showing that reversible assembly/disassembly is a major mechanism of regulating V-ATPase activity and the luminal pH of endolysosomes/lysosomes in mammalian cells (Trombetta *et al*., 2003; Lafourcade *et al*., 2008; Stransky and Forgac, 2015; Ratto *et al*., 2022). Our data, from continuously fed NRK cells, contrast, but are not incompatible, with previous reports of changes in nutrient status, specifically amino acid availability, resulting in mTORC1 regulation of V-ATPase assembly, such that when mTORC1 activity declines, the V1 sub-complex assembles with the Vo sub-complex to form functional protein pumps to acidify lysosomal organelles(Ratto *et al*., 2022). The finding that V-ATPase disassembly during ELR is not dependent on mTOR activation is consistent with a recent report suggesting that lysosome reformation in ELR in fed cells uses a mechanism that does not require mTOR activation to provide reformed lysosomes(Rodgers *et al*., 2022). This mechanism, suggested to require a phosphoinositide pathway, differs from that underlying ALR, which has been shown to require activation of mTORC1 in cells subjected to prolonged serum starvation (Yu *et al*., 2010).

Analysis of events occurring during ELR has proved challenging because budding, tubulation and scission can occur rapidly and are not synchronised throughout the cell (Bright *et al*., 2005; Bissig *et al*., 2017). Recently, an approach to circumventing these problems has been developed using spinning disc microscopy and a rapid imaging/analysis workflow to quantitate events(Rodgers *et al*., 2022). Our approach was different, taking advantage of our previous observation that the generation of sucrosomes in NRK cells does not activate autophagy or result in TFEB translocation to the nucleus(Bright *et al*., 2016) unlike in many other cell types (Sardiello *et al*., 2009; Higuchi *et al*., 2015). It provided us with cells in which all terminal lysosomes had been recruited into swollen endolysosomes and from which ELR could be activated approximately synchronously following uptake of invertase.

Our experiments investigating the colocalization of GFP-tagged proteins with LAMP1-mCherry by confocal microscopy, suggested that the V1-ATPase protein V1G1 was recruited to the limiting membrane of sucrosomes during their formation, along with RAB7a and RILP and dissociated with a similar time course to RAB7a and RILP when lysosomes reformed after invertase uptake. This was in contrast to the integral membrane protein Voa3 and also ARL8b, whose colocalization with LAMP1-mCherry did not alter significantly. That the changes in colocalization of GFP-tagged V1G1 represented reversible assembly and disassembly of the whole V-ATPase during sucrosome formation and lysosome reformation was confirmed by the immunoblotting of magnetically enriched lysosome and sucrosome preparations (LysoPrep and SucroPrep), including those made following invertase uptake. Further evidence of increased V-ATPase assembly during endolysosome formation was obtained from the live cell experiments demonstrating the recruitment of cytosolic V1G1 to the endolysosome limiting membrane soon after the commencement of luminal content mixing during endosome-lysosome kissing and fusion.

Our findings about the colocalization of RAB7a and ARL8b with each other and with LAMP1, during sucrosome formation and lysosome reformation, are consistent with previous reports in other cell types of reduced RAB7a density and an enrichment of ARL8b relative to RAB7a on peripheral lysosomes compared to juxtanuclear lysosomes (Johnson *et al*., 2016)as well as the proposal that the ARL8b effector SKIP together with HOPS recruits the RAB7a GAP TBC1D15 resulting in what has been termed RAB7a to ARL8b conversion with microtubule plus-end mediated transport of organelles to the cell periphery (Jongsma *et al*., 2020). Our colocalization data do not distinguish between either the increased colocalization of RAB7a with LAMP1 after formation of sucrosomes being simply a reflection of the recruitment of peripheral RAB7a-depleted/LAMP1-enriched lysosomes into the swollen sucrosomes as a result of fusion events with RAB7a-positive late endosomes or the direct recruitment from a cytosolic pool onto the sucrosomes, although we did not investigate this further. Nevertheless, the data support a model in which increased RAB7a on the endolysosomal membrane is involved in the recruitment of the V1 subcomplex and loss of RAB7a is linked to reversible V-ATPase disassembly during ELR, in continuously fed cells. Exactly where and when disassembly occurs during lysosome reformation is unclear. We observed that GFP-tagged V1G1 and RAB7a are present in the reformation tubules extending from sucrosomes after invertase uptake, but very depleted on the reformed, less acidic, peripheral lysosomes. This suggests that loss of the V1 ATPase subcomplex occurs after scission of the reformation tubules at a stage equivalent to the protolysosome in ALR (Yu *et al*., 2010) (Lenardo). However, scission itself is not absolutely necessary for loss of the V1 subcomplex, because we showed that the dynamin inhibitor dynasore, which had a profound morphological effect consistent with preventing scission, had no effect on the reduction in colocalization of GFP-tagged V1G1 with LAMP1-mCherry after invertase uptake into sucrosomes.

Our data suggest that it is unlikely that the previously well documented binding of the RAB7a effector RILP to the VIG1 subunit of the V-ATPase (De Luca *et al*., 2014) is sufficient to explain fully the reversible assembly/disassembly of the V-ATPase. The published observation that over-expression of the dominant negative T22N mutant of RAB7a resulted in strongly reduced accumulation of the acidotropic probe LysoTracker Red, provided compelling evidence that RAB7a is directly involved in regulating endolysosomal/lysosomal pH (Bucci *et al*., 2000). However, for RILP, the situation is more confused with reports of RILP depletion causing increased organelle acidification (lysotracker intensity) (De Luca *et al*., 2014), having little or no effect (cresyl violet accumulation)(Ratto *et al*., 2022) or causing alkalinization (ratiometric imaging) (Johnson *et al*., 2016). We saw no effect of RILP depletion on pH in NRK cells stably expressing the lysosomal pH probe pHLARE, on the increase of colocalization of EGFP-tagged V1G1with LAMP1-mCherry during formation of sucrosomes or on its decrease following invertase uptake. Further complexity is suggested by evidence that RILP remains bound to V1G1 in cells expressing the T22N mutant of RAB7a, but is no longer recruited to lysosomal membranes and that RILP also promotes VIG1 degradation (De Luca and Bucci, 2014; De Luca *et al*., 2014).

It has been proposed that mTORC1-regulation of lysosomal acidification is mediated through reversible, DMXL1-dependent and not RILP-dependent, recruitment of the V1 subcomplex to lysosomal membranes (Ratto *et al*., 2022). In experiments supporting this proposal, genetic ablation of DMXL1 strongly suppressed lysosomal V1B2 levels as well as accumulation of the acidotropic dye cresyl violet under basal as well as mTORC1-inhibited conditions, consistent with it also being involved in V-ATPase assembly in untreated, continuously fed cells. Mammalian DMXL1 and WDR7 (also known as raboconnectin 3α and 3β) are related to components of the yeast V-ATPase assembly factor known as the RAVE complex (Jaskolka *et al*., 2021). Intriguingly, both these proteins have been implicated as RAB7a interactors by proximity biotinylation. They were identified as ‘strong hits’ using proximity biotinylation with a mitochondrially-localised form (MitoID) of constitutively active (Q67L) RAB7a (Gillingham *et al*., 2019) and were also identified as potential interactors of both wild type and constitutively active RAB7a, with greatly reduced interaction with dominant negative (T22N) RAB7a, using BioID (data set S1 in(Yan *et al*., 2022)). Thus, given the dynamic equilibrium observed between V1 and Vo subunits, implied from the rapid recovery times in our FRAP experiments, it may be that the loss of RAB7A during lysosome reformation, and thereby the inability to recruit DMXL1/WDR7, is sufficient to result in net disassembly of the V-ATPase, although we have not tested this directly. We also cannot rule out the possible involvement of a variety of other reported factors that bind V-ATPase and have been suggested to regulate its assembly, disassembly, stability and/or activity, including several members of the TLDc (TBC (Tre-2/Bub2/Cdc16) LysM domain catalytic) family of proteins(Eaton *et al*., 2021; Tan *et al*., 2022; Wang *et al*., 2022; Wilkens *et al*., 2023), lysosomal STK11IP (Zi *et al*., 2022) and the cytosolic chaperonin tailless complex polypeptide 1 ring complex (TRiC)(Merkulova *et al*., 2015; Ratto *et al*., 2022).

The lack of change in colocalization of ARL8b with LAMP1 throughout the lysosome fusion and regeneration cycle compared with the increase in colocalization of RAB7a with LAMP1 during sucrosome (swollen endolysosome) formation and decrease during ELR, raises a number of questions that require future investigation. These include whether RAB7a is recruited to the sucrosomes from the cytosol, despite this seeming unlikely because, as has been pointed out in a different context (Schleinitz *et al*., 2023), the presence of ARL8b on organelles, should counteract the recruitment of RAB7a, through ARL8b’s attraction of the RAB7a GAP TBC1D15. Given that we did not see loss of RAB7a and disassembly of the V-ATPase until after scission of lysosome reformation tubules, except when scission was inhibited, other important questions include the exact timing of TBC1D15 recruitment, RAB7a loss and V-ATPase disassembly. A further task for future work is to explore the mechanisms of tubule formation and fission during lysosome reformation in mTORC1-independent ELR, which may have both similarities(Boutry *et al*., 2023) and differences from that in mTORC1-regulated ALR and has been suggested to require the sorting nexin SNX2(Rodgers *et al*., 2022). Based on our findings to date, a model of V-ATPase reversible assembly/disassembly throughout the lysosome fusion and regeneration cycle in continuously fed cells is presented in Fig 7.

**Figure 7.**
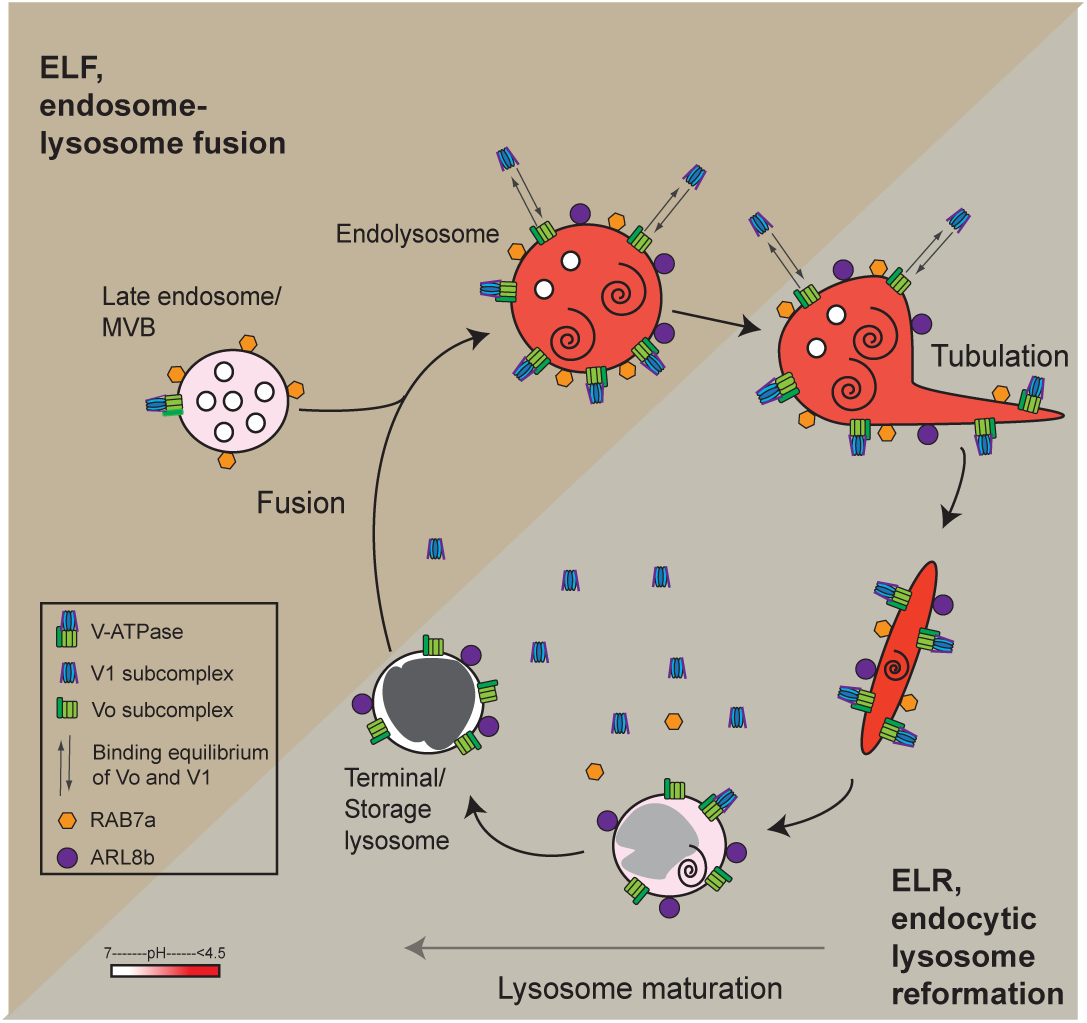
V-ATPase assembly and disassembly during the lysosome regeneration cycle in continuously fed cells. Scheme showing the net assembly of V-ATPase on the limiting membrane of endolysosomes formed by fusion of terminal storage lysosomes with late endosomes and net disassembly during reformation of lysosomes. The dynamic equilibrium of the V1 and Vo subcomplexes on the endolysosome membrane is also shown as well as the presence/absence of RAB7a and ARL8b. Tubulation, maturation, and content condensation processes required for the reformation of terminal storage lysosomes are indicated. Endolysosomes can undertake further fusions with late endosomes and/or terminal storage lysosomes. Acidic compartments are shaded pink (slightly acidic) to dark red (pH <4.5). Electron-dense content is shaded dark gray.

## Materials and Methods

### Reagents

Dulbecco’s modified Eagle’s medium (DMEM), Roswell Park Memorial Institute medium (RPMI 1640), fetal calf serum (FCS), Dulbecco’s phosphate buffered saline (PBS), glutamine, penicillin, streptomycin, hygromycin B, HEPES,TES, cOmplete^™^ Protease Inhibitor Cocktail, poly L-lysine (MW 30-70 kDa) and invertase were from Merck Life Science UK Ltd, Gillingham, Dorset, UK. Magic Red^™^ Cathepsin B was from ImmunoChemistry Technologies (Bloomington, MN USA) and BMV109 from Vergent Biosciences, Minneapolis, MN USA. Nigericin (sodium salt) and monensin (sodium salt) were from the Cayman Chemical Company (Ann Arbor, MI, USA). Geneticin (G418), Dextran-Alexa 546, Dextran-Alexa 647 and Dextran-Oregon Green 488 (all three MW 10,000, anionic, fixable), CellTracker™ Blue CMAC dye, AEBSF protease inhibitor, latrunculin A, DTT and DMEM without Phenol Red were from ThermoFisher Scientific (Waltham, MA, USA). 10 000 MW DexoMAG^TM^ was from Liquids Research Ltd (Bangor, Gwynedd, UK). Rabbit polyclonal antibodies to LAMP1 (to a cytosolic tail epitope, ab24170), GFP (ab6556) and mouse IgG (ab6709); rabbit monoclonal antibodies to calreticulin (ab92516), EEA1 (ab109110), Na^+^/K^+^ ATPase (ab76020), TOMM20 (ab186735) and Vod1 (ab202899); donkey anti-rabbit Ig Alexa 488 (ab150073), Alexa 594 (ab150064) and Alexa 680 (ab175772); donkey anti-mouse IgG Alexa 488 (ab150105, Alexa 594 (ab150108) and Alexa 790 (ab175780) were from Abcam (Cambridge, UK). Rabbit polyclonal antibodies to TGN38 and cation independent mannose 6-phosphate receptor (ciMPR) were as previously described (Reaves *et al*., 1996). Mouse monoclonal anti vinculin (V9131) and rabbit polyclonal anti beta actin (A2066) were from Merck. Rabbit polyclonal antibodies to ARL8b (A88749), mCherry(A85306) and Voa3 (for immunoblotting, A91446) were from Antibodies.com, Cambridge, UK; V1G1 (#16143) and V1A1 (#17115) from Proteintech Europe, Manchester, UK; to RILP (NBP1-98279) from Novus Biologicals/Bio-techne Ltd Abingdon, UK; to phospho-4E-BP1 (#9459), phosphor-p70 S6 kinase (#9234) phospho-S6 ribosomal protein (#2217) from Cell Signaling Technology Inc, Danvers, MA, USA. Rabbit monoclonal antibody to RAB7a (#D95F2) was from Cell Signaling Technology. Mouse monoclonal antibodies to GAPDH (am4300) and GFP (ma5-15256) were from Invitrogen/Thermofisher Scientific and to Golg2a/GM130 (#66662) from Proteintech.

The affinity purified rabbit polyclonal antibodies to human VPS18 (aa 40-394)(Graham *et al*., 2013) and to Voa3 (peptide PDASTLENSWSPDEEK)(Lange *et al*., 2006) (a gift from Dr Thomas Jentsch, Max-Delbrück-Centrum für Molekulare Medizin, Berlin, Germany), were as previously described. Protein A gold (PAG) conjugates were from the Department of Cell Biology, University of Utrecht. Nigericin (sodium salt) and monensin (sodium salt) were from the Cayman Chemical Company (Ann Arbor, MI, USA). Torin 1 was from Tocris Bioscience (Bristol, UK), dynasore monohydrate from Santa Cruz Biotechnology (Dallas, TX, USA).

### Cell Culture, Transduction and siRNA knockdown

NRK fibroblasts were cultured in a 5% CO_2_ humidified incubator at 37°C in DMEM supplemented with 10% (v/v) FCS, 100 IU/ml penicillin, 100 μg/mL streptomycin, 4.5 g/L glucose, and 2 mM L-glutamine (this supplemented DMEM referred to as DMEM below unless stated) and Phoenix cells cultured in RPMI similarly supplemented, but without antibiotics. Cells were negative for mycoplasma contamination (routinely tested for the presence of mycoplasma contamination using MycoAlert™ Mycoplasma Detection Kit (LT07-318) from Lonza Bioscience Blackley, UK, and treated with mycoplasma removing agent BM-Cyclin (#10 799 050 001) from Roche Diagnostics, Burgess Hill, UK). To create stably expressing NRK cells (Supplementary Table S1), EGFP and mCherry tagged cDNA constructs were cloned into pBMN and modified pLXIN (Clontech, Mountain View, CA, USA) retroviral expression vectors(Peden *et al*., 2004; Gordon *et al*., 2009). pHLARE(Webb *et al*., 2021) (sfGFP-LAMP1-mCherry, Addgene plasmid # 164477, a gift from Diane Barber), was subcloned into pLXIN. Retroviral infections were performed as described previously(Peden *et al*., 2004; Gordon *et al*., 2009)after packaging with the Phoenix cell packaging system(Swift *et al*., 2001). The transduced NRK cells were selected with either 0.5mg/mL geneticin (G418) (pLXIN) or 0.2 mg/mL hygromycin B (pBMN). Clonal lines were established after cell sorting on a BD Influx5 cell sorter. For transduced NRK cells expressing GFP-tagged human RILP, endogenous rat RILP was depleted with a Dharmacon^TM^ ON-TARGET plus siRNA SMARTpool (#L-084959-02) from Horizon Discovery Biosciences Ltd, Cambridge, UK using a double transfection knockdown protocol described previously(Parkinson *et al*., 2015).

For microscopy, cells were cultured after seeding onto 13 mm or 25mm glass coverslips for live- or fixed-cell confocal microscopy, Deckgläser 25 mm coverslips (Techmate Ltd, Milton Keynes, UK) for ratiometric imaging, glass bottom dishes ± grids (MatTek Ashland, MA, USA) for time-lapse live cell imaging, CLEM and EM of cell monolayers unroofed by coverslip rip-off or 25 cm^2^ tissue culture flasks for TEM or immuno-EM.

### Formation of sucrosomes / reformation of lysosomes

To form sucrosomes in the late endocytic pathway, cells were cultured in DMEM containing 30 mM sucrose for 24h at 37°C(Bright *et al*., 1997). In experiments in which we investigated lysosome reformation including the extrusion of reformation tubules from sucrosomes, the sucrose-containing DMEM was replaced with DMEM containing 0.5 mg/mL invertase at 37°C for the times indicated in individual experiments.

### Endocytic Compartment Labelling with Fluorescent Dextran and Cathepsin Activity Substrates

Unless stated, terminal endocytic compartments in cultured NRK fibroblasts were loaded with 0.5 mg/ml lysine-fixable Dextran-Oregon Green or dextran-Alexa 647 in DMEM for 4 h at 37°C followed by incubation in conjugate-free DMEM for 20 h as previously described(Bright *et al*., 2005). Early endocytic compartments were loaded with dextran-Alexa 546 for 10 min followed by a 5 min chase in conjugate-free medium. To label endocytic organelles in which cathepsins were catalytically active, cells were incubated at 37°C with 673 nM cathepsin B MR (Magic Red™) substrate in DMEM without Phenol Red for at least 2 min as previously described(Bright *et al*., 2016) or 1μM BMV109 for 30 min (Repnik *et al*., 2017).

### Subcellular fractionation and magnetic isolation of lysosomes

To prepare membrane and cytosol fractions, NRK cells were seeded onto 10cm dishes, allowed to reach ∼75% confluence, rinsed twice with ice-cold phosphate buffered saline, pH 7.4 (PBS) and fractions prepared at 4°C. The cells were scraped into 650µL of homogenization buffer (250mM sucrose, 1 mM EDTA, 10 mM HEPES pH 7.4 containing cOmplete™ Protease Inhibitor Cocktail at concentration recommended by supplier), lysed by 20 strokes with an Isobiotec ball-bearing cell homogenizer using the 8µm tungsten carbide ball and after removal of a nuclear pellet (centrifugation 7000g, 5min), the cell lysate was centrifuged at 100,000g for 30 min to pellet the membrane fraction. Membrane pellets were rinsed twice with homogenization buffer and resuspended in 100µL homogenization buffer containing 1% SDS. The 100,000g supernatant (the cytosolic fraction) was concentrated to 100µL with an Amicon Ultra 10K centrifugal filter from Merck according to manufacturer’s instructions.

For the magnetic isolation of lysosomes and sucrosomes, untreated NRK cells or cells containing sucrosomes, at ∼75% confluence on 15 cm dishes, were incubated for 1h in DMEM containing 1mg/mL 10 000 MW DexoMAG^TM^, followed by a 4h chase in DMEM alone. For some cells containing sucrosomes, the chase (4h or 8h) was in DMEM containing 0.5 mg/mL invertase. Rinsing, homogenization and generation of a post-7000g cell lysate was as above, except that the homogenization buffer (1 mL per dish) used was STM (250mM sucrose, 1 mM MgCl_2_, 10 mM TES pH 7.4 containing cOmplete™ Protease Inhibitor Cocktail at concentration recommended by supplier). Isolation of the lysosomes was then as previously described(Rofe and Pryor, 2016), but with final elution from the MiniMACS^TM^ Large Cell Column (Miltenyi BioTec Ltd, Bisley, UK) into 150μL STM.

Protein concentration of subcellular fractions was measured with a Pierce^TM^ bicinchonic acid assay kit (ThermoFisher) using bovine serum albumin (BSA) to generate a standard curve.

### Immunoblotting

Immunoblotting was as previously described(Davis *et al*., 2021) except that SDS-PAGE was carried out using Millipore mPAGE® 10% Bis-Tris Precast Gels (MP10W10/ MP10W12/ MP10W15) with MES SDS running buffer (MPMES) from Merck, before protein transfer to PVDF membranes.

### Microscopy

Preparation, fixation and labelling of cells for fluorescence and electron microscopy, preparation of thin sections for transmission EM (TEM) and immuno-EM of frozen sections, use of microscopes, image analysis and quantification were carried out as previously described(Bright *et al*., 2016), unless stated. Confocal and live cell microscopy were carried out on Zeiss 710 and 780 and 880 confocal microscopes (CarlZeiss, Ltd., Welwyn Garden City, UK) and EM sections were examined with an FEI Tecnai G2 Spirit BioTwin TEM (Eindhoven, The Netherlands).

### Quantification of fluorescence microscopy and visualisation/calculation of organelle pH

Fluorescence colocalization analysis including calculation of Pearson’s and Manders’ colocalization coefficients was carried out using open source CellProfiler software (https://cellprofiler.org/), using its automated puncta detection and thresholding pipeline on confocal images. Usually, three independent experiments were performed with images of 10 randomly selected cells taken per condition. Unless stated, colocalization was measured in whole cells. For some experiments, colocalization was estimated in an outer cytoplasmic shell after cells were incubated with 10µM CellTracker™ Blue for 15min prior to imaging. CellTracker™ Blue positive individual cell images were each detected as a primary object in CellProfiler and the object shrunk by 20μm from the cell edge to obtain the inner shell (the perinuclear region). This was subtracted from the original object to obtain the outer shell (the peripheral region).

Visualisation of individual organelle pH was achieved, after endocytic uptake of fluorescent dextrans, by dual-fluorochrome imaging utilising Oregon Green 488 as the pH-sensitive fluorochrome and Alexa 647 as the pH-insensitive fluorochrome(Bright *et al*., 2016). Quantification of organelles pH was by measuring, using CellProfiler, the ratio of fluorescence intensity of mCherry and sfGFP of individual organelles in cells stably expressing pHLARE(Webb *et al*., 2021). Individual organelles of different pH were automatically placed in bins corresponding to pH values <4.5, 4.5-4.9, 5-5.4, 5.5-5.9, 6-6.5 and >6.5 using the Classify Objects function of CellProfiler and also a mean pH value for the organelles in a whole cell calculated. A pH standard curve was constructed by incubating the cells in different pH buffers in the presence of nigericin and monensin (pH clamped cells) as previously described(Bright *et al*., 2016).

### Fluorescence Recovery after Photobleaching

Stable NRK cell lines expressing Voa3-EGFP or V1G1-EGFP grown on MatTek glass bottom dishes were incubated with 30 mM sucrose for 24h, transferred to the 37°C stage of an LSM780 confocal microscope and incubated with Magic Red™ as above to label the catalytically active sucrosomes. Live cells were imaged using a x63 1.4 NA Plan Apochromat oil-immersion lens using an Argon laser line at 488 nm (used to excite EGFP) and a Diode laser line at 561 nm (used to excite cresyl violet). Laser power was attenuated to 2% of maximum to minimize photobleaching and phototoxicity. The detector pinholes were set to give a 0.5-0.9 μm optical slice. Pixel dwell times were 1.27μs using multitracking (line switching) with a line average of 4. Photobleaching of the EGFP in a region of interest (ROI) was achieved using 20 iterations of the 488 nm laser line at 100% intensity and post-bleach images were collected at 5-15s intervals to determine recovery of EGFP fluorescence into the bleached region. Cresyl violet fluorescence was used to identify the bleached organelle when an entire sucrosome was selected as the ROI. Acquisition and quantitation of the fluorescence intensity of the limiting membrane or tubules was performed using Zen software (Zen Black 2.3 SP1). Line scans were used to determine the peak membrane intensity at 8 points and the mean value recorded for each timepoint of the FRAP time-lapse series. The data was incorporated into GraphPad Prism (v 5.01) for graphical representation and movies were assembled using NIH Image J (v2.0.0-rc-69/1.52).

### Cell unroofing and pre-embedding immunolabelling for EM

For pre-embedding immunolabelling of samples for EM, NRK cells were grown on MatTek glass bottom dishes. The medium was aspirated and the cells were washed with stabilization buffer [30 mM HEPES pH 7.4, 70 mM KCl, 5 mM MgCl_2_, 3 mM EGTA, 1 mM DTT and 0.1 mM AEBSF protease inhibitor] for 1 min at 37°C. A coverslip coated with 1 mg/ml poly L-lysine was then placed over the well of the MatTek dish and the stabilisation buffer aspirated to allow the coverslip to come to rest upon the cell monolayer. The cells were then unroofed by coverslip rip-off(Miller *et al*., 1991) [S7] and immediately fixed with 2% paraformaldehyde (PFA) in PBS at 37°C. The unroofed cells were washed with PBS, unreacted aldehydes were quenched with 50 mM NH_4_Cl for 10 min at 20°C and blocked with 1% BSA / 5% FCS in PBS for 10 mins at 20°C. Pre-embedding immunolabelling was then performed with mouse or rabbit primary antibodies. Rabbit anti-mouse IgG was used as a bridging antibody and the primary or bridging rabbit antibodies were detected by adding PAG. The samples were then washed with PBS and 0.1M Na cacodylate Buffer, pH7.2 and processed for TEM.

### Statistical Methods

Data are presented ± SEM of means of separate experiments. The statistical significance of variations between means for different conditions was determined using the GraphPad Prism v 5.01 statistical package, applying the non-parametric Kruskal-Wallis test and Dunn’s post hoc test.

## Supporting information

Supplemental Sava et al Reversible assembly

## Acknowledgements

This work was supported by the MRC (research grant MR/R0009015/1 to J.P.L. and N.A.B), and the BBSRC with GSK Research and Development Ltd [industrial CASE studentship to I.S. and L.J.D]. We are indebted to Matthew Gratian and Mark Bowen from the CIMR core microscopy facility for technical assistance and thank Geoffrey Hesketh and David Gershlick for helpful discussions and comments on the manuscript.

## Abbreviations

ALR: autophagic lysosome reformation
ELF: endosome-lysosome fusion
ELR: endocytic lysosome reformation.

## Videos

**Video 1.** Time lapse confocal fluorescence microscopy movie of V1G1-EGFP entering lysosome reformation tubule after incubating V1G1-EGFP cells containing sucrosomes with 0.5 mg/mL invertase for 1h in the presence of Magic Red cathepsin B substrate.

**Video 2.** Time lapse confocal fluorescence microscopy movie of an NRK cell containing sucrosomes incubated for 2h with 0.5 mg/mL invertase, the second hour + 80 μM dynasore in the presence of Magic Red cathepsin B substrate.

V1G1-EGFP entering lysosome reformation tubule after incubating V1G1-EGFP cells containing sucrosomes with 0.5 mg/mL invertase for 1h in the presence of Magic Red cathepsin B substrate.

**Video 3.** Time lapse confocal fluorescence microscopy movie of a clonal V1G1-EGFP cell showing kissing and fusion of an individual Dex A546-positive late endosome (red) with Dex A647-positive terminal endocytic compartments (lysosomes) resulting in the acquisition of Dex A647 (blue) in the Dex A546-positive organelle and the rise in recruitment of V1G1-EGFP (green) to the limiting membrane of the resulting endolysosome.

**Video 4.** Time lapse confocal fluorescence microscopy movies of FRAP of individual sucrosomes of clonal NRK cells expressing V1G1-EGFP or Voa3-EGFP. An intensity profile is shown below indicating high-intensity (red), mid-intensity (green) or low-intensity (blue) fluorescence.

