## Supplemental Sava et al Reversible assembly for "Reversible assembly and disassembly of V-ATPase during the lysosome regeneration cycle"

**Supplemental Material**

Supplemental Table S1

and Supplemental Figures S1 -S6

**Supplemental Table 1**

**Table S1 Stable expression of tagged proteins in transduced NRK cells**

| Stably expressing cells (mixed stable, ms or clone, #) | Vector | Protein immuno-blotted | Overexpression (fold) |
|---|---|---|---|
| pHLARE ms | pLXIN | LAMP1 | 0.77 |
| V1G1-EGFP ms | pLXIN | V1G1 | 8.7 |
| V1G1-EGFP #7 | pLXIN | V1G1 | 7.1 |
| Voa3-EGFP #1 | pLXIN | Voa3 | 2.4 |
| LAMP1-EGFP#10 | pLXIN | LAMP1 | 12.5 |
| LAMP1-mCherry#6 | pBMN | LAMP1 | 1.3 |
| LAMP1-mCherry#6/Voa3-EGFP ms | /pLXIN | Voa3 | 1.7 |
| LAMP1-mCherry#6/V1G1-EGFP ms | /pLXIN | V1G1 | 7.8 |
| LAMP1-mCherry#6/ RAB7a-EGFP ms | /pLXIN | RAB7a | 0.2 |
| LAMP1-mCherry#6/ EGFP-RILP #4 | /pLXIN | RILP | 1.4 |
| LAMP1-mCherry#6/ ARL8b ms | /pLXIN | ARL8b | 0.8 |
| ARL8b-mCherry/RAB7a-EGFP | pBMN/pLXIN | ARL8b<br>RAB7a | 0.8<br>0.7 |

Overexpression was determined by quantitative immunoblotting and expressed as fold change relative to endogenous protein in control NRK cells. LAMP1 was rat LAMP1 also known as lgp120. LAMP1-mCherry#6 cells were used to create cells co-expressing LAMP1-mCherry and various EGFP-tagged proteins as listed. The pHLARE pH probe is based on rat LAMP1 and expression was compared with endogenous LAMP1.

### Figure S1

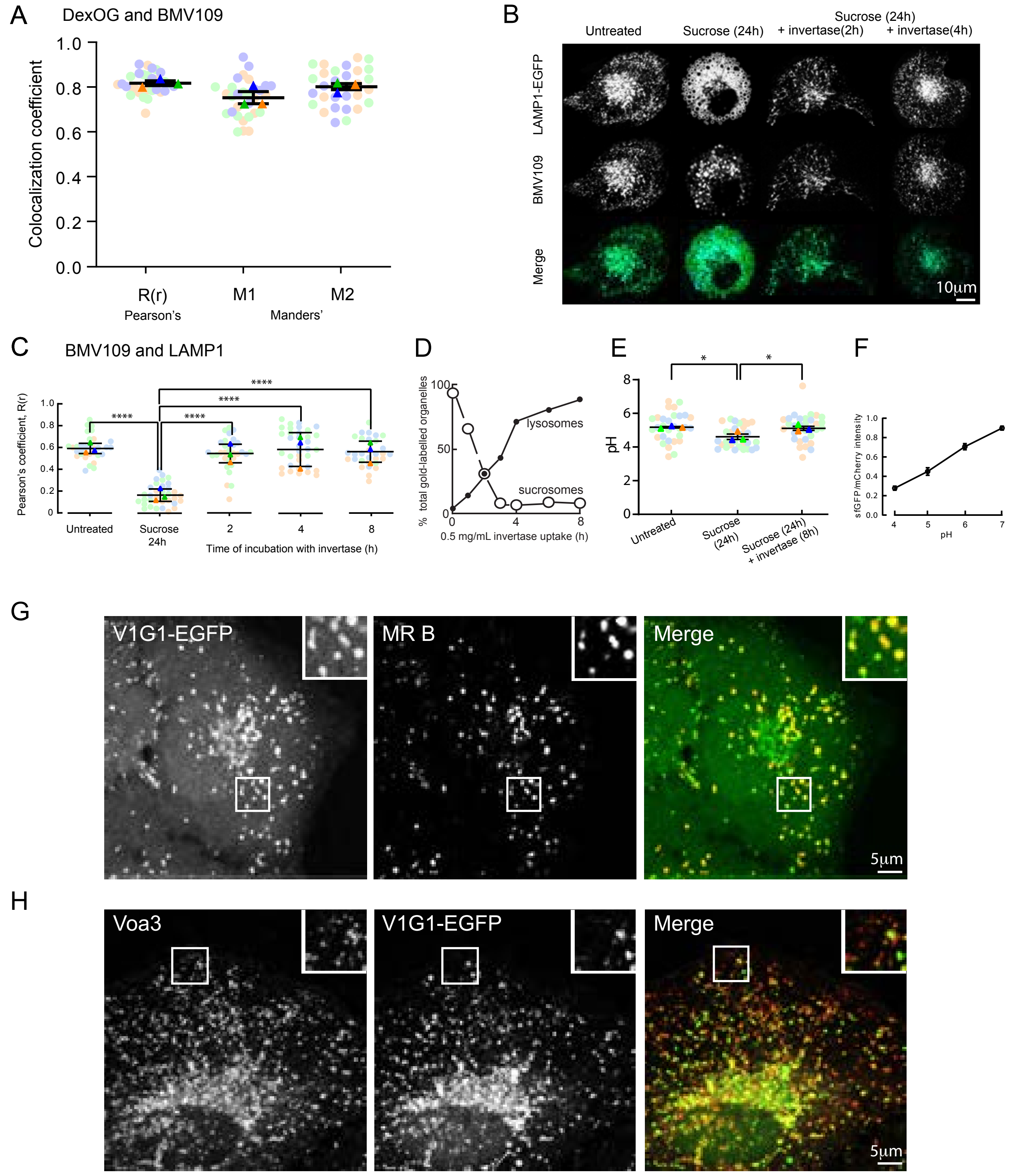


**Legend on next page**

**Figure S1. Cathepsin activity, pH and V-ATPase subunit localization in NRK cells. (A)** Pearson's (R(r),Mander's (M1, DexOG:BMV109) and M2, BMV109:DexOG) correlation coefficients (mean ± SEM of 3 experiments, 10 cells each) for colocalization of pre-loaded DexOG (4h uptake/20h chase) and BMV109 (30min). Representative images from single cell shown in Fig 1B. **(B)** Confocal fluorescence microscopy images of representative NRK cells stably expressing LAMP1-EGFP, otherwise untreated, incubated with 30mM sucrose for 24h or incubated with sucrose followed by 2h or 4h incubation with invertase, all incubated with BMV109 for 30min. **(C)** Pearson's correlation coefficients (mean ± SEM of 3 experiments, ≥10 cells each) for colocalization of LAMP1-EGFP and BMV109 in live NRK cells stably expressing LAMP1-EGFP, otherwise untreated, incubated with 30mM sucrose for 24h or incubated with sucrose followed by 2, 4 or 8h incubation with invertase, all incubated with BMV109 for 30min. **(D)** Time course of loss of sucrosomes and appearance of dense core lysosomes re-plotted from previously published EM data(Bright *et al.*, 1997), which quantitated the distribution of pre-loaded BSA-gold (4h uptake, 20h chase) in NRK cells after incubation with 30mM sucrose for 24h followed by incibation with 0.5 mg/mL invertase. **(E)** pH of pHLARE-positive organelles in NRK cells stably expressing pHLARE and otherwise untreated, incubated with 30mM sucrose for 24h or incubated with sucrose followed by 8h incubation with 0.5mg/mL invertase. Mean ± SEM of 3 experiments in each of which the mean pH of all pHLARE-positive organelles in each of 10 cells (single confocal section of each cell) was calculated by comparison with a standard curve shown in panel (F). *, $p < 0.05$. **(F)** pH standard curve derived from pH clamped NRK cells cells stably expressing pHLARE. Mean ±SEM of 3 experiments (10 cells per condition) shown**. (G)** Confocal fluorescence images of a live cell expressing V1G1-EGFP (from mixed stables) incubated with Magic Red cathepsin B substrate (MR B) for 2min prior to imaging. Confocal fluorescence images of a fixed cell expressing V1G1-EGFP (from mixed stables) labelled with an antibody to Voa3(Lange *et al.*, 2006) prior to imaging.

### Figure S2

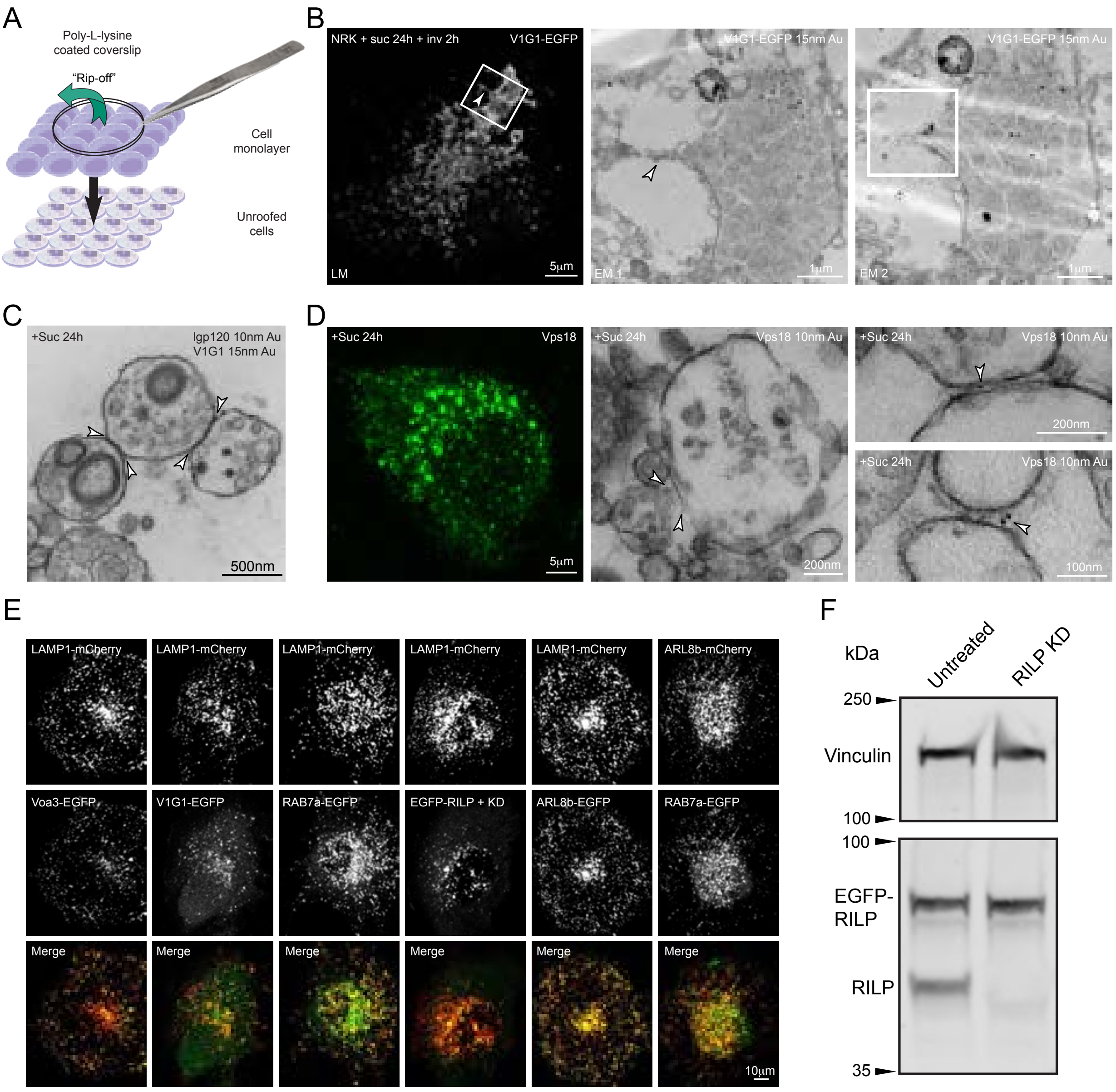


**Legend on next page**

**Figure S2. EM unroofing, CLEM, images of cells expressing mCherry-tagged proteins together with various EGFP-tagged proteins and RILP depletion. (A)** Schematic diagram showing technique for unroofing of cultured cells prior to pre-embedding labelling for immunoEM. **(B)** CLEM showing confocal immunofluorescence image of an unroofed NRK cell expressing V1G1-EGFP, which had been incubated with 30mM sucrose for 24h and 0.5mg/mL invertase for 2h and was labelled after unroofing with anti-GFP, alongside EM images of two serial sections of the boxed area. The right-hand EM image has a box highlighting the area shown in Fig 2E. **(C)** Immunoelectron micrograph of part of an unroofed NRK cell expressing V1G1-EGFP, which had been incubated with 30mM sucrose for 24h and was labelled after unroofing with antibodies to rat LAMP1 (lgp120) and GFP (V1G1). Arrowheads show edges of regions of close apposition of the sucrosomes with filamentous tethers linking the organelles. **(D)** Confocal immunofluorescence image of an unroofed NRK cell which had been incubated with 30mM sucrose for 24h and was labelled after unroofing with anti-VPS18, alongside representative immunoEM images of tethered sucrosomes showing sparse VPS18 labelling at the edges and occasionally in the tethered regions. **(E)** Representative confocal fluorescence microscopy images of NRK cells stably expressing LAMP1-mCherry or ARL8b-mCherry and various EGFP-tagged proteins. Cells were mixed stables other than the clonal LAMP1-mCherry/EGFP-RILP cell which was from a population also depleted of endogenous RILP by siRNA-mediated knockdown. **(F)** Representative immunoblot of cell lysates from untreated LAMP1-mCherry/EGFP-RILP cells and cells after siRNA-mediated knockdown (KD) of endogenous RILP.

### Figure S3

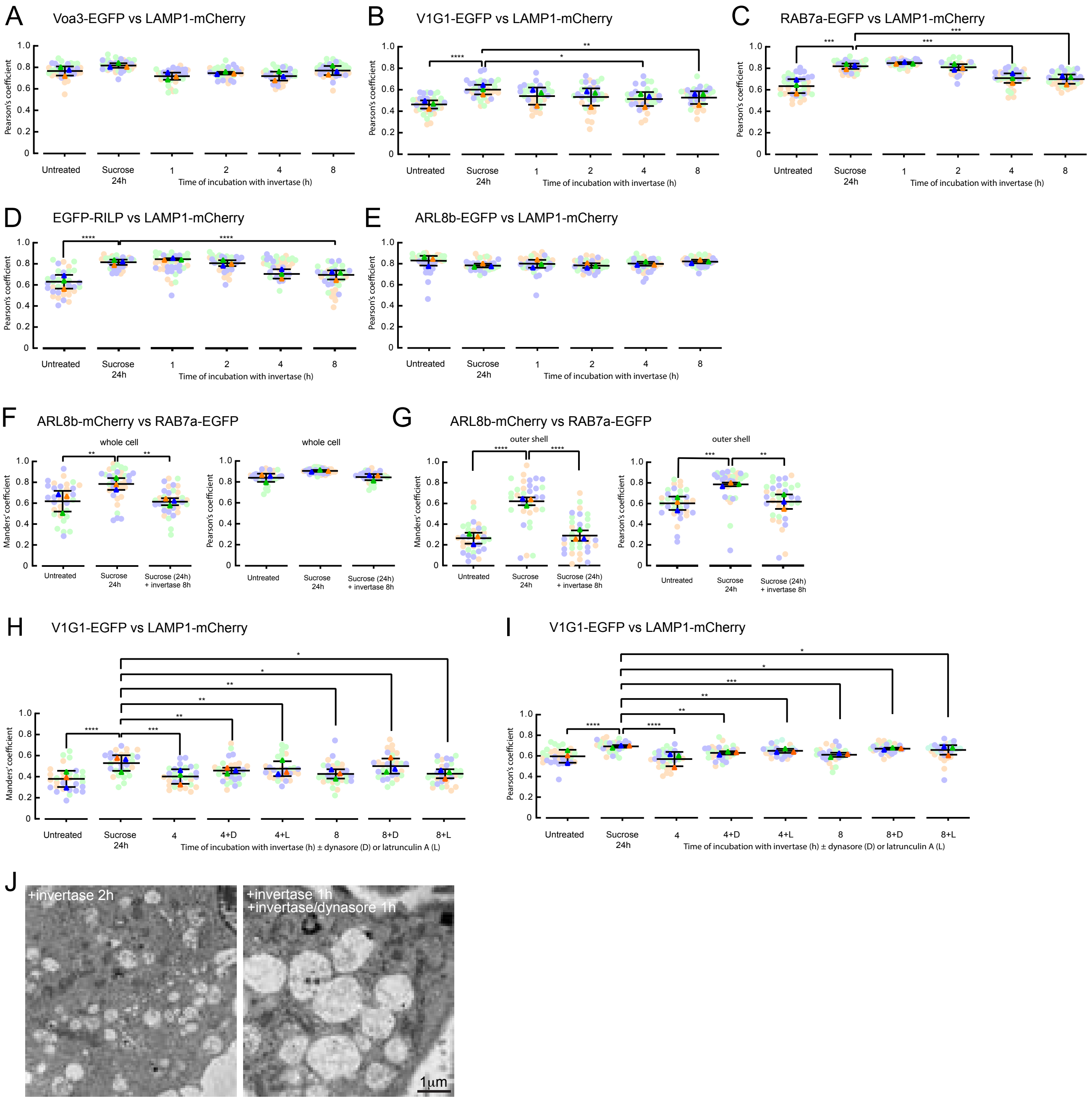


**Legend on next page**

**Figure S3. Colocalization of EGFP-tagged proteins with LAMP1-mCherry or ARL8b-mCherry in stably expressing NRK cells. (A-E)** Pearson's colocalization coefficients for EGFP-tagged proteins vs LAMP1-mCherry in untreated cells, cells incubated with 30mM sucrose for 24h or incubated with sucrose followed by 1-8h incubation with 0.5mg/mL invertase. Mixed stable cells used except in (D) in which a clonal LAMP1-mCherry/ RILP-EGFP cell line depleted of endogenous RILP by siRNA-mediated knockdown (86.7±1.9% (3) depletion) was used. Pearson's colocalization coefficients are shown for exactly the same experiments as the Manders' colocalization coefficients shown in Fig 3 A-E. **(F, G)** Manders' and Pearson's colocalization coefficients (whole cell, F; outer shell, G) for ARL8b-mCherry vs RAB7a-EGFP in untreated mixed stable ARL8b-mCherry/ RAB7a-EGFP cells, cells incubated with 30mM sucrose for 24h or incubated with sucrose followed by 8h incubation with 0.5mg/mL invertase. The individual data points are for the same cells shown in Fig 3G. **(H, I)** Manders'(H) colocalization coefficient for V1G1-EGFP vs LAMP1-mCherry and Pearson's (I) colocalization coefficient in the presence or absence of 80μM dynasore or 1μM latrunculin A in otherwise untreated LAMP1-mCherry/ V1G1-EGFP mixed stable cells, cells incubated with 30mM sucrose for 24h or incubated with sucrose followed by 8h incubation with 0.5mg/mL invertase. The individual data points are for the same cells shown in Fig 3I. **(J)** Electron micrographs of portions of NRK cells incubated with 30mM sucrose for 24h followed by 2h incubation with 0.5mg/mL invertase ± 80μM dynasore during the second h, showing the appearance of 'daisy chains' of linked swollen sucrosomes after dynasore treatment. In panels A-I, mean ± SEM of 3 experiments (minimum 10 cells per experiment) shown. *, $p<0.05$; **, $p<0.01$, ***, $p<0.005$; ****, $p<0.001$.

#### Figure S4

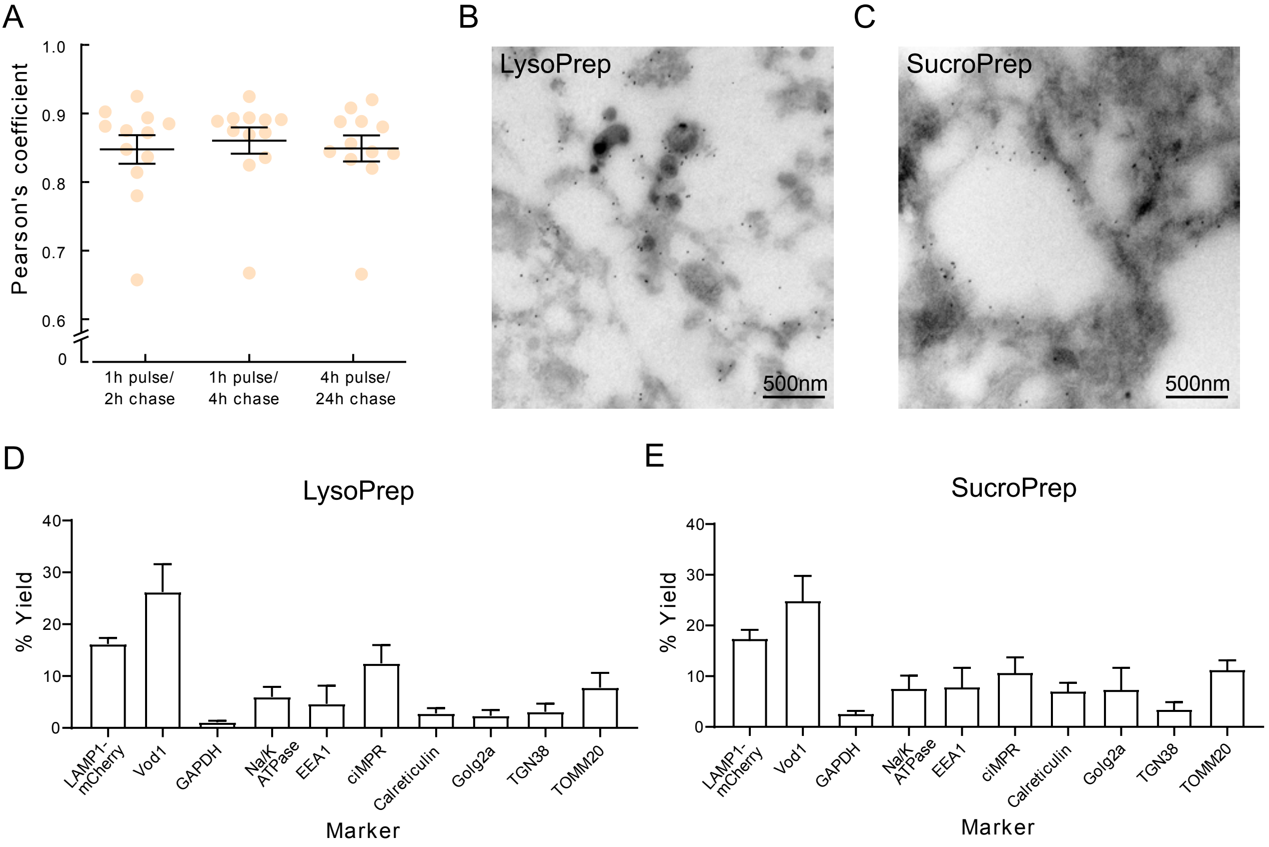


**Figure S4. Characterization of LysoPrep and SucroPrep. (A)** Terminal endocytic compartments of NRK cells were pre-loaded by incubating the cells with Dex A647 for 4h followed by a 20h chase in Dex A-free medium, and then the cells were incubated with DexOG for 1h followed by a 2 or 4h chase in Dex OG-free medium or 4h followed by a 20h chase in Dex OG-free medium. Pearson's colocalization coefficients of 12 cells from each condition in a single experiment shown and with no significant difference observed in the mean colocalization coefficient (mean±SEM shown). **(B, C)** Immunoelectron micrographs of magnetically isolated lysosomes (LysoPrep, B) and sucrosomes (SucroPrep, C) from LAMP1-mCherry/V1G1-EGFP cells showing anti-LAMP1 labelling (gold particles), on dense core lysosomes in LysoPrep and at the periphery of sucrosomes in Sucroprep. **(D, E)** Yield (recovery relative to lysate shown as mean±SEM of data from 4 experiments) of various subcellular markers calculated from immunoblots (for representative example see Fig 4E) of Lysopreps and Sucropreps from LAMP1-mCherry/V1G1-EGFP cells.

**Figure S5**

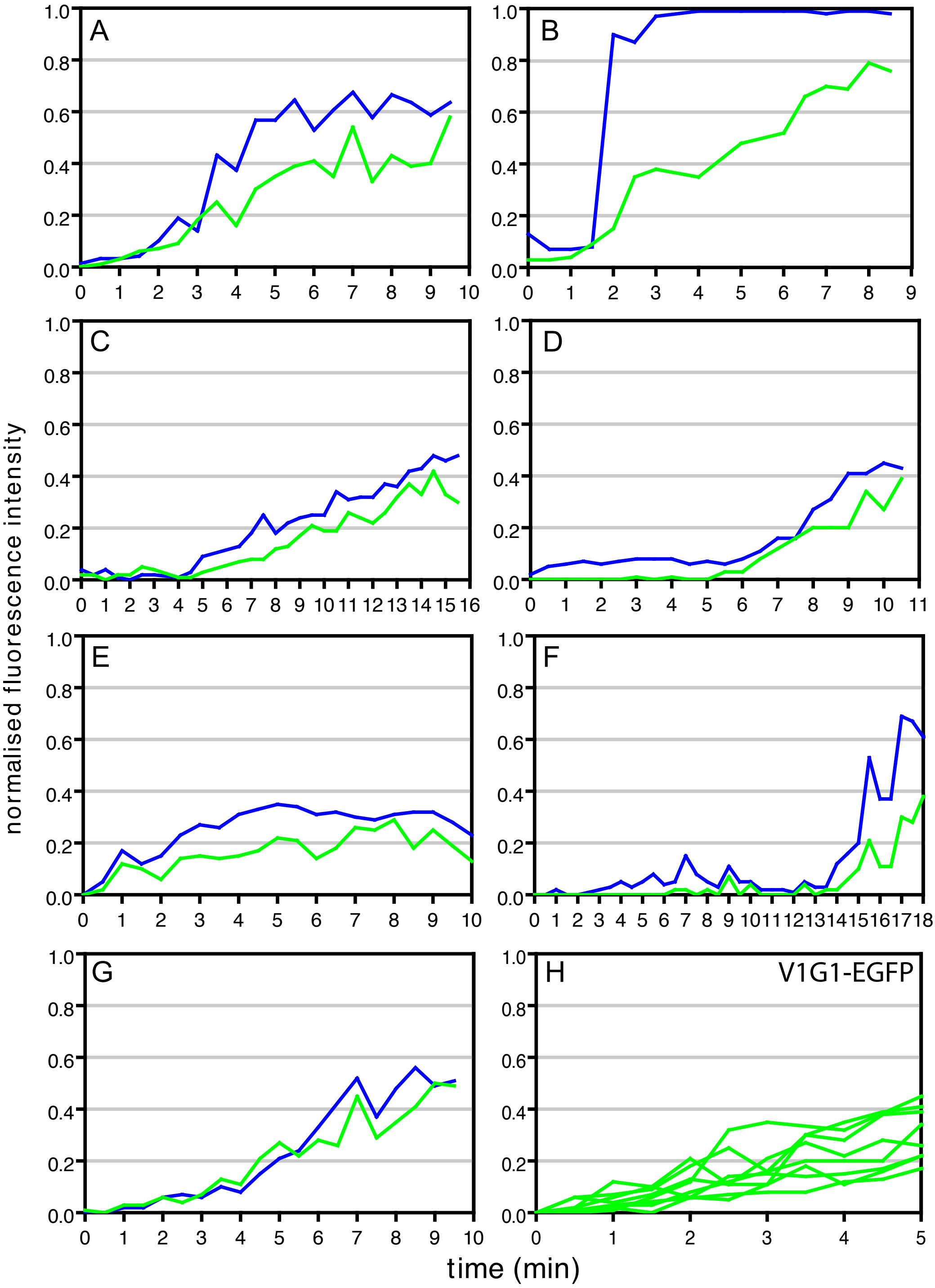


**Legend on next page**

**Figure S5. The dynamics of V-ATPase V1 subunit recruitment to endolysosomes in live cells. (A-G).** Further examples to that shown in Fig 5 A, B of kissing and fusion of individual Dex A546-positive late endosomes with Dex A647-positive terminal endocytic compartments (lysosomes) resulted in the gradual acquisition of Dex A647 (blue lines) in the Dex A546-positive organelle and the rise in recruitment of V1G1-EGFP (green) to the limiting membrane of the resulting endolysosome. Clonal NRK cells expressing V1G1-EGFP were treated and examined exactly as in the Fig5A legend and mean grayscale intensity profiles of Dex A647 (blue) and V1G1-EGFP (green) plotted over time for each Dex A546-positive organelle undergoing kissing and/or fusion that was examined. **(H)** Summary of the time course of increase in V1G1-EGFP fluorescence intensity on the limiting membrane of the organelles shown in panels A-G from the point when Dex A647 begins to increase.

### Figure S6

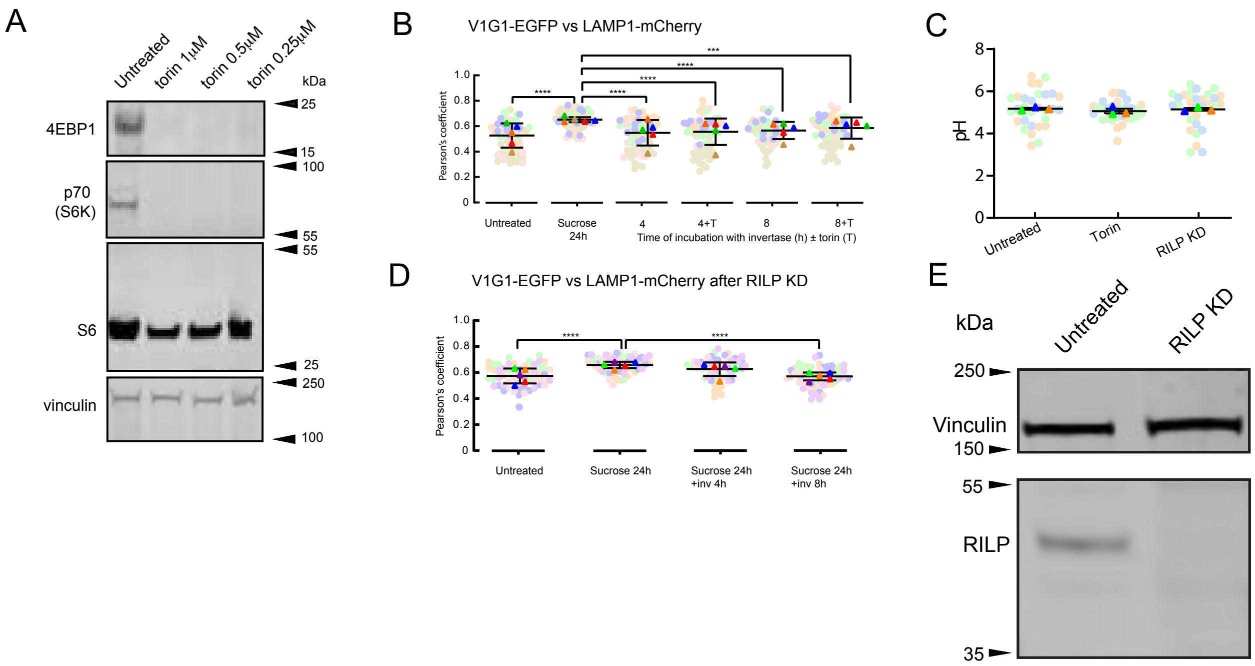


**Figure S6. Effects of torin and RILP depletion on V-ATPase assembly/ disassembly and organelle pH. (A)** Immunoblot of phosphorylated mTORC1 substrates in NRK cells showing that three concentrations of torin (0.25, 0.5, 1 µM) applied for 15min reduce phosphorylation to the same extent. **(B)** Pearson's colocalization coefficients for V1G1-EGFP vs LAMP1-mCherry in otherwise untreated LAMP1-mCherry/ V1G1-EGFP mixed stable cells, cells incubated with 30mM sucrose for 24h or incubated with sucrose followed by 4 or 8h incubation with 0.5mg/mL invertase ± 0.25µM torin. Mean ± SEM of 3 experiments (minimum 10 cells per experiment) shown. ***, p<0.005; ****, p<0.001. The individual data points are for the same cells shown in Fig 6C. **(C)** pH of pHLARE-positive organelles in NRK cells stably expressing pHLARE and otherwise untreated, incubated with0.25  M torin 15min or incubated with sucrose or depleted of RILP (96.1±1.3% (3) depletion) by siRNA-mediated knockdown. Mean ± SEM of 3 experiments in each of which the mean pH of all pHLARE-positive organelles in each of 10 cells (single confocal section of each cell) was calculated by comparison with a standard curve shown in Fig S1F. **(D)** Pearson's colocalization coefficients for V1G1-EGFP vs LAMP1-mCherry in LAMP1-mCherry/ V1G1-EGFP mixed stable cells following siRNA-mediated knockdown of RILP (85.1±2% (5) depletion) and otherwise untreated, incubated with 30mM sucrose for 24h or incubated with sucrose followed by 4 or 8h incubation with 0.5mg/mL invertase. Mean ± SEM of 5 experiments (minimum 10 cells per experiment) shown. ****, p<0.001. The individual data points are for the same cells shown in Fig 6G. **(E)** Representative immunoblot of cell lysates from untreated LAMP1-mCherry/ V1G1-EGFP mixed stable cells and cells after siRNA-mediated knockdown (KD) of RILP.
